# Semantic and syntactic composition of minimal adjective-noun phrases in Dutch: an MEG study

**DOI:** 10.1101/2020.03.14.991802

**Authors:** Arnold Kochari, Ashley Lewis, Jan-Mathijs Schoffelen, Herbert Schriefers

## Abstract

The possibility to combine smaller units of meaning (e.g., words) to create new and more complex meanings (e.g., phrases and sentences) is a fundamental feature of human language. In the present project, we investigated how the brain supports the semantic and syntactic composition of two-word adjective-noun phrases in Dutch, using magnetoencephalography (MEG). The present investigation followed up on previous studies reporting a composition effect in the left anterior temporal lobe (LATL) when comparing neural activity at nouns combined with adjectives, as opposed to nouns in a non-compositional context. The first aim of the present study was to investigate whether this effect, as well as its modulation by noun specificity and adjective class, can also be observed in Dutch. A second aim was to investigate to what extent these effects may be driven by syntactic composition rather than primarily by semantic composition as was previously proposed. To this end, a novel condition was administered in which participants saw nouns combined with pseudowords lacking meaning but agreeing with the nouns in terms of grammatical gender, as real adjectives would. We failed to observe a composition effect or its modulation in both a confirmatory analysis (focused on the cortical region and time-window where it has previously been reported) and in exploratory analyses (where we tested multiple regions and an extended potential time-window of the effect). A syntactically driven composition effect was also not observed in our data. We do, however, successfully observe an independent, previously reported effect on single word processing in our data, confirming that our MEG data processing pipeline does meaningfully capture language processing activity by the brain. The failure to observe the composition effect in LATL is surprising given that it has been previously reported in multiple studies. Reviewing all previous studies investigating this effect, we propose that materials and a task involving imagery might be necessary for this effect to be observed. In addition, we identified substantial variability in the regions of interest analysed in previous studies, which warrants additional checks of robustness of the effect. Further research should identify limits and conditions under which this effect can be observed. The failure to observe specifically a syntactic composition effect in such minimal phrases is less surprising given that it has not been previously reported in MEG data.

## Introduction

One of the fundamental properties of human language is compositionality - the possibility to combine smaller units of meaning into larger ones creating new, integrated meanings. This is possible at multiple levels; for example, morphemes are combined into words, words are combined into phrases, phrases are combined into sentences. Here, we focus on how our brains combine individual words into phrases - for example, when ‘large’ and ‘insect’ are combined into ‘large insect’. To achieve that, our brain has to retrieve the meaning of each word and combine them in terms of the meaning (semantic composition), but also in terms of structural dependency, i.e., which word is a modifier and which word is modified (syntactic composition).

Different approaches have been taken in cognitive neuroscience to unpack the processes underlying such composition. Some studies looked into the composition of meaningful phrases and sentences as opposed to ones with a semantic or a syntactic violation (for example, lines of work looking at N400, LAN and P600 event-related potential signatures; Kaan, Harris, Gibson, & Holcomb, 2000; Kutas & Federmeier, 2011; Lau, Phillips, & Poeppel, 2008) or as opposed to unstructured word lists (e.g., measuring fMRI BOLD; Friederici, Meyer, & von Cramon, 2000; Hultén, Schoffelen, Uddén, Lam, & Hagoort, 2019; Humphries, Binder, Medler, & Liebenthal, 2006). Other studies manipulated the level of semantic or syntactic complexity of a sentence or a phrase (e.g., Bornkessel, Zysset, Friederici, von Cramon, & Schlesewsky, 2005; Makuuchi, Bahlmann, Anwander, & Friederici, 2009; Pallier, Devauchelle, & Dehaene, 2011). However, processing a longer phrase or sentence necessarily involves some processes in addition to composition such as storage of information in working memory, ambiguity resolution, pragmatic or discourse inferences etc. One approach that avoids these potential additional processes is looking at composition in minimal two-word phrases (‘large insect’) as compared to processing a single word (‘insect’). In this case, the composition is stripped to the most basic process: as opposed to processing a single word, the only added processes should be retrieval of the meaning of the additional word (adjective) and composition. While natural language utterances are clearly more complex, we can take composition of such a minimal phrase as a starting point for investigating brain dynamics supporting linguistic combinatory processing.

In the present study, we investigate semantic and syntactic composition of two-word phrases as compared to processing a single word in Dutch using magnetoencephalography (MEG).

### Composition of minimal adjective-noun phrases: MEG studies

A series of MEG studies by Pylkkänen and colleagues investigated spatial and temporal aspects of processing minimal adjective-noun phrases as opposed to processing nouns preceded by consonant strings (e.g., Bemis & Pylkkänen, 2011, 2013a, 2013b; Del Prato & Pylkkänen, 2014; Flick et al., 2018; Pylkkänen, Bemis, & Blanco Elorrieta, 2014; Westerlund & Pylkkänen, 2014; Ziegler & Pylkkänen, 2016 etc.; see Pylkkänen, 2016 for a review). In these studies, participants were presented with either an adjective and a noun (e.g. ‘red car’ - composition condition) or a meaningless consonant string and a noun (e.g. ‘xgf car’ - no composition condition; a consonant string equalizes the amount of visual input to the brain in both conditions) in a word-by-word fashion. Subsequently, they were presented with a picture or a written question and judged whether it corresponded to the meaning of the phrase or the noun, depending on the condition. These studies consistently report higher levels of activity in the left anterior temporal lobe (LATL) in the composition condition as opposed to the no composition condition approximately 200-250 ms after noun onset (henceforth, we will refer to this difference as the ‘LATL composition effect’). Similar composition-related activity has been reported for the processing of noun-noun compounds (Flick et al., 2018; Zhang & Pylkkänen, 2015) and minimalistic verb phrases (Kim & Pylkkänen, 2019; Westerlund, Kastner, Al Kaabi, & Pylkkänen, 2015). Besides English, involvement of LATL in composition of minimalistic two-word phrases in a similar time-window has also been reported for Modern Standard Arabic (Westerlund et al., 2015) and American Sign Language (Blanco-Elorrieta, Kastner, Emmorey, & Pylkkänen, 2018). Altogether, these results were interpreted as demonstrating that LATL is the brain region most distinctly responsible for composition of the meaning of two words (Pylkkänen, 2016; Pylkkänen & Brennan, 2019). It should be noted, however, that the localization and timing of the LATL composition effect varied considerably between studies (reviewed in *Table 5, General Discussion*). It was sometimes located in areas beyond what would typically be considered the anterior portion of the left temporal lobe. The earliest onset and the latest offset time of the effect varied between roughly 180 ms and 350 ms, lasting for approximately 50-100 ms. The presence of such variability was one of the motivations for conducting the present study. We discuss these points in detail below.

In the above-mentioned series of MEG studies, composition-related brain activity has also been reported in several other regions, in addition to LATL, though not consistently: the right anterior temporal lobe in approximately the same time-window as LATL but also in a later time window (Bemis & Pylkkänen, 2011; Poortman & Pylkkänen, 2016), ventromedial prefrontal cortex at approximately 350-500 ms after noun onset (Bemis & Pylkkänen, 2011; Del Prato & Pylkkänen, 2014) and left angular gyrus (at approximately 350-400 ms after noun onset in the visual modality and 540-590 ms after noun onset in the auditory modality; Bemis & Pylkkänen, 2013a). Given that increased activity in these regions was not observed consistently across studies, these effects have to be considered as less robust than the effect in LATL. To anticipate, for these reasons we will, in the present study, focus primarily on the LATL effect when looking at the contrast between the composition and no composition conditions described above.

### Investigating semantic composition of minimal phrases

On a theoretical level, one can distinguish between semantic and syntactic composition processes. The contrast between an adjective-noun phrase and a consonant string-noun combination employed in the above described MEG studies does not allow one to distinguish between these two processes as both are involved in the former and both are absent in the latter. However, because the LATL composition effect appears to be modulated by semantic-conceptual factors (discussed in detail below), it has largely been interpreted as reflecting semantic (or conceptual) composition rather than syntactic composition Pylkkänen, 2016; Pylkkänen & Brennan, 2019; Westerlund & Pylkkänen, 2014).

Besides the MEG studies, several fMRI studies investigated which brain regions show increased levels of activity for semantic composition processes in minimal two-word phrases. In one such study, Price and colleagues (Price, Bonner, Peelle, & Grossman, 2015) presented participants with adjective-noun phrases differing in plausibility (i.e., how meaningful the phrase was - e.g., ‘loud car’ vs #‘moss pony’). The processing of more meaningful phrases was accompanied by a higher BOLD signal in both the left and right angular gyri (BA39^1^) than the processing of less meaningful phrases. This difference was interpreted as evidence for angular gyrus playing a crucial role in semantic composition (see also Price, Peelle, Bonner, Grossman, & Hamilton, 2016, for supporting causal evidence from transcranial direct current stimulation). Another study (Matchin, Hammerly, & Lau, 2017) contrasted processing determiner-noun phrases and determiner-pseudoword phrases. Here, the authors reasoned that whereas in the case of determiner-noun phrases semantic composition should take place, such semantic composition would be greatly reduced for the determiner-pseudoword phrases since pseudowords do not carry any associated semantic meaning. More BOLD activity was observed for processing determiner-noun phrases in a small region in the posterior portion of the middle temporal gyrus (BA21), presumably showing that this region is more involved in semantic composition. Yet another study (Schell, Zaccarella, & Friederici, 2017) compared brain activity in adjective-noun phrases (e.g. ‘blaues Schiff’ [blue ship]), determiner-noun phrases (e.g. ‘dieses Schiff’ [this ship]), and single nouns (e.g. ‘Schiff’ [ship]). Schell and colleagues viewed the adjective-noun composition as ‘semantically driven’ and reasoned that the regions more involved in processing of these phrases as opposed to determiner-noun phrases (which they view as ‘syntactically driven’ since they have reduced descriptive content) and single nouns should be partaking in semantic composition^2^. Processing the adjective-noun phrases in comparison to single nouns was accompanied by more BOLD activity in the left inferior frontal gyrus (LIFG; BA45), and in comparison to determiner-noun phrases it was accompanied by more BOLD activity in the left angular gyrus (BA39). The regions identified in all of these studies can be seen as additional potential candidates for carrying out semantic composition in minimal adjective-noun phrases: left and right angular gyri, left posterior temporal lobe, and left inferior frontal gyrus.

To investigate semantic composition of two-word phrases further, in this project we will primarily focus on whether we can observe the LATL composition effect, as reported in the MEG studies discussed above. In addition, we will investigate whether the semantic composition effect might be present in the regions indicated by the fMRI studies, though we will do the latter only in an exploratory manner since the regions in which effects were observed in the fMRI studies did not seem to exhibit differential activity in the MEG studies.

### Modulation of the LATL composition effect by noun specificity

In an attempt to look further into the specific process that the LATL composition effect reflects, several MEG studies varied the type of stimuli within the experimental set-up outlined above. One factor in this context is noun specificity (Westerlund & Pylkkänen, 2014; Zhang & Pylkkänen, 2015; Ziegler & Pylkkänen, 2016). In these studies, no composition effect (i.e., no difference between adjective-noun and consonant string-noun combinations) was observed in LATL when the noun in the adjective-noun phrase denoted a more specific meaning (e.g. *rose, trout*), whereas the composition effect *was* present when the noun denoted a less specific meaning (e.g., *flower, fish*; Westerlund & Pylkkänen, 2014; Ziegler & Pylkkänen, 2016). These findings were interpreted as a reflection of a narrowing down of the meaning of the noun: when the noun has a rather specific meaning, the information provided by an adjective provides relatively little additional information concerning potential referents. By contrast, when the noun is less specific, the adjective adds more information with respect to potential referents. The fact that a conceptual feature of the noun appears to modulate the composition effect suggests that the effect reflects semantic composition rather than syntactic composition since phrases with high and low specific nouns do not differ syntactically (Pylkkänen, 2016; Pylkkänen & Brennan, 2019). However, the modulation of the composition effect was observed in somewhat differing regions of interest (within or close to LATL) in the three relevant studies (specifically, left BA38 in Zhang & Pylkkänen, 2015; left BA21 in Ziegler & Pylkkänen, 2016; left BA38+21+20 in Westerlund & Pylkkänen, 2014). Moreover, one of the studies that looked at noun specificity reported finding a confounding difference in adjective-noun plausibility (i.e., typicality) between high and low specificity conditions in a post-hoc norming study that they conducted after their data was collected (Westerlund & Pylkkänen, 2014), whereas the other two do not report controlling for plausibility (Zhang & Pylkkänen, 2015; Ziegler & Pylkkänen, 2016). Thus, though a clear modulation of the LATL composition effect by noun specificity would be important for the interpretation of this effect, it appears that this modulation needs additional investigation to establish its robustness and stability, and concerning potential confounds.

### Modulation of the LATL composition effect by adjective class

Differences between types of adjectives might also have a modulating role, and might thus allow for a further specification of the composition processes reflected in the LATL composition effect (Ziegler & Pylkkänen, 2016). The meaning of one class of adjectives, so called *scalar adjectives*, is strongly context dependent - their meaning largely depends on the noun that they are combined with. For example, ‘large’ refers to different sizes when combined with ‘fruit’, ‘horse’ or ‘house’. In theoretical semantics, this is captured by the assumption that the noun determines the comparison class for the property denoted by the adjective. For example, in the case of ‘large’ the comparison class would need to consist of typical sizes of either fruits, horses or houses (e.g., Kennedy, 2007; Kennedy & McNally, 2005; Klein, 1980; van Rooij, 2011). These scalar adjectives are also sometimes called *gradable* or *vague*; other examples are ‘tall’, ‘long’, ‘loud’, ‘heavy’ etc. By contrast to scalar adjectives, the meaning of so-called *intersective adjectives*^3^ does not depend on context, or does so to a much lesser degree. The meaning of an intersective adjective such as ‘square’, ‘dead’ or ‘ceramic’ remains relatively stable for different nouns (naturally, some ambiguity always remains, but we assume that this ambiguity is drastically smaller for intersective compared to scalar adjectives).

Ziegler and Pylkkänen (2016) contrasted adjective-noun combinations comprised of scalar adjectives or intersective adjectives and found more activity in the time-window of the LATL composition effect – 200-300 ms after noun onset – for nouns combined with intersective adjectives than for nouns combined with scalar adjectives (though the ROI was again somewhat different from other studies on the LATL composition effect). Ziegler and Pylkkänen interpret this difference as suggesting that composition of nouns with intersective adjectives happens at the standard time-window of the LATL composition effect, whereas composition of the noun with scalar adjectives does not or cannot yet happen at this time-window, but happens later. Specifically, they propose this interpretation assuming that the activation of semantic features of a noun happens in a gradient fashion, with retrieval proceeding from general to highly specific features (Moss, McCormick, & Tyler, 1997; Pylkkänen, 2016). They further assume that composition of a noun with a scalar adjective requires for the noun meaning to be rather specific since scalar adjectives’ meaning depends on a specific comparison class, whereas composition of a noun with intersective adjectives does not require the same degree of specificity because the adjective meaning depends less on the comparison class of the noun. Hence, when participants see a noun preceded by an intersective adjective, composition can happen early, when still relatively little information about the noun is retrieved (at 200-300 ms), whereas when they see a scalar adjective, more information about this noun has to be retrieved before composition can happen, so it happens later (therefore, no composition effect at 200-300 ms yet).

However, the Ziegler and Pylkkänen study is the only one to date that reports the modulation of the LATL composition effect by the adjective class, and also with a region of interest slightly different from the original studies reporting the LATL composition effect. In addition, as in the studies on the role of noun specificity described above, also in this study on adjective types, there was no explicit control for matching the plausibility of the adjective-noun phrases between conditions. Thus, as for the potential effect of noun specificity, it appears that the modulation of the composition effect by adjective class also should be checked for robustness and stability.

### Investigating syntactic composition of minimal phrases

In contrast to semantic composition, syntactic composition has received much less attention in MEG studies, but has been investigated in a number of fMRI studies. Here, we discuss research on syntactic composition with a focus on two-word phrases.

As already mentioned, the experimental design of the MEG studies on the LATL composition effect cannot directly distinguish between semantic and syntactic composition effects. In the respective conditions contrasted in these studies, either both aspects of composition (semantic and syntactic) were simultaneously present or simultaneously absent. Thus, LATL is also a candidate region for performing syntactic composition in minimal phrases. Moreover, a number of fMRI studies have shown that LATL might be sensitive to manipulations of syntactic information (e.g., Brennan et al., 2012; Rogalsky & Hickok, 2009; Schell et al., 2017), and this even holds for sentences consisting exclusively of pseudowords where lexico-semantic information is fully absent (e.g., Humphries et al., 2006; Mazoyer et al., 1993; however, see Flick & Pylkkänen, 2018; Pylkkänen & Brennan, 2019 for arguments on why sentences with pseudowords cannot be considered completely void of meaning). On the other hand, it should be noted that the importance of LATL for syntactic processing has been questioned; for example, patients with an atrophy in the LATL do not show signs of syntactic processing deficit (Wilson et al., 2014; see Matchin & Hickok, 2020 for this argument and other arguments against LATL being important for syntactic processing).

Several recent fMRI studies focused specifically on identifying brain regions responsible for syntactic composition in minimal phrases. Zaccarella and Friederici (2015) investigated brain activity when processing a two-word phrase consisting of a determiner and a pseudoword in German (e.g., ‘DIESE FLIRK’ [this flirk]). Syntactically, such a phrase is clearly a determiner phrase, but it has no meaning (i.e., no semantic component). Compared to a two-word list condition (e.g. ‘APFEL FLIRK’ [apple flirk]) which lacked syntactic structure, the processing of determiner-pseudoword phrases resulted in more BOLD activity in a portion of the left inferior frontal gyrus (BA44). In a similar study that used real words instead of pseudowords, Zaccarella and colleagues (Zaccarella, Meyer, Makuuchi, & Friederici, 2017) observed an increase in activity related to syntactic composition again in LIFG (BA44, BA45), but also in left posterior superior temporal sulcus (BA22). Consistent with these results, the already mentioned study of Schell and colleagues (Schell et al., 2017) reported more BOLD activity for processing determiner phrases in comparison to single word processing and in comparison to adjective-noun phrase processing also in LIFG (BA44, BA45; but also BA47) and in left posterior superior temporal sulcus and middle temporal gyrus (BA21, BA22). Based on these studies, the LIFG as well as parts of the posterior temporal lobe potentially play a crucial role in syntactic composition processes. Note, however, that not all fMRI studies on syntactic composition observed these effects: in another fMRI study that looked at minimal phrase composition, Matchin and colleagues (Matchin et al., 2017) did not observe any differences between processing a two-word phrase and a non-structured list; we will return to this point in the *General Discussion*.

To the best of our knowledge, there is until now no MEG study looking specifically at syntactic composition processes for minimal adjective-noun phrase processing. In the present study, we will therefore include a condition targeting specifically syntactic composition processes in addition to conditions targeting semantic (and syntactic) composition.

### Present study

In the present study, we will primarily look into composition processes as reflected by the LATL composition effect reported in previous studies, and, more specifically, into potential modulation of the LATL composition effect by semantic-conceptual properties of the adjective and the noun in adjective-noun phrases, which support its interpretation as reflecting specifically semantic composition. In addition, we will look into syntactic composition processes in adjective-noun phrases. We do so using MEG which will allow us to identify corresponding activity with high temporal resolution and to identify the brain regions that are involved.

The literature review provided above shows that while multiple MEG studies reported a composition effect in LATL (starting from Bemis & Pylkkänen, 2011), these previous findings are mixed in terms of spatial and temporal extent of the effect. In the present study, we will use a design that is parallel to the one used in the MEG studies by Pylkkänen and colleagues. Within this general design, we will compare the processing of adjective-noun phrases with the processing of single nouns (i.e., nouns preceded by a letter string); the contrast between these two conditions should reflect the composition processes in phrases. We will focus on the LATL composition effect, i.e., more activity in LATL at approximately 200-250 ms after noun onset for the adjective-noun phrase as opposed to single nouns. In addition, we will introduce two factors that appear to modulate composition processes and are, thus, important for understanding the observed effect: noun specificity (see Westerlund & Pylkkänen, 2014; Zhang & Pylkkänen, 2015; Ziegler & Pylkkänen, 2016) and adjective class (see Ziegler & Pylkkänen, 2016).

In the present study we chose to closely follow the design and analysis choices of a previous study. Specifically, we closely follow the study by Ziegler and Pylkkanen (2016; henceforth referred to as Z&P) since it included both the noun specificity and the adjective class manipulations that we are interested in. In our analyses, we compared neural activity of processing nouns preceded by real adjectives as opposed to preceded by a letter strings (i.e., single noun processing). We used the same region of interest and time-windows in which the Z&P study reported their observed effects as our pre-specified hypotheses in confirmatory analyses. However, as there is some inconsistency in the localization and timing of the reported composition effect in previous studies, we also planned exploratory analyses of other regions and time-windows where composition effects have previously been reported (see *Method* for details on different analyses that we conducted).

The second goal of the present study was to investigate syntactic composition processes in adjective-noun phrases within the same set-up as for semantic composition. The involved brain regions and the time-course of syntactic composition with such minimalistic phrases has not yet been addressed in MEG studies. To this end, we included a condition in which instead of an adjective the participants saw pseudowords combined with real nouns (similar to a number of studies that used such ‘jabberwocky’ stimuli in the past - e.g., Matchin et al., 2017; Mazoyer et al., 1993; Pallier et al., 2011; Zaccarella & Friederici, 2015). These pseudowords were inflected for grammatical gender like real adjectives and are, therefore, henceforth referred to as ‘pseudoadjectives’. This was possible because the present study was conducted in Dutch (rather than English, a language where the following described properties are not present). In indefinite noun phrases with a prenominal adjective in Dutch, the adjective agrees with the noun in terms of the noun’s grammatical gender - it either has an ending ‘e’ or does not depending on which gender the noun belongs to (e.g., ‘klein paard’[small horse], ‘kleine tafel’ [small table] but *‘kleine paard’, *‘klein tafel’). The pseudoadjectives in our study agreed with nouns in terms of the grammatical gender, i.e., there was morphosyntactic agreement. We expected that such pseudoadjective-noun phrases would induce syntactic composition in the absence of semantic composition (see *Materials* below for a detailed discussion of a control for potential semantic associations). Given that no previous published MEG studies targeted specifically syntactic composition processes in minimal phrases, we did not have strong hypotheses about the time-course and regions where they should be observable. We thus compared the neural activity of processing nouns preceded by pseudoadjectives as opposed to nouns preceded by letter strings (i.e., single noun processing) in an exploratory way in the regions that have been implicated for syntactic composition by fMRI studies investigating minimal phrases.

## Method

### Participants

This study fell under the ethical approval for standard studies at Donders Institute for Brain, Cognition and Behaviour, Radboud University Nijmegen, by ‘Commissie Mensgebonden Onderzoek Regio Arnhem-Nijmegen’. The data was collected between February and May 2019. The number of participants, criteria for participation and inclusion in the analyses were set before the start of the data collection. To take part in the experiment, the participants had to be native speakers of Dutch, 18-35 years old, without any language-related impairment (such as dyslexia), had to have normal or corrected-to-normal vision, and should not have taken part in any of the studies where the experimental stimuli were pre-tested (described below). In addition, the participants had to meet MEG/MRI inclusion criteria at the Donders Institute (specifically, not claustrophobic, no metal in or on the body, no dental wire, no pacemaker etc.). Forty participants were recruited for this study in exchange for a payment according to the local regulations.

To be included in the analysis, participants had to reach overall accuracy on comprehension questions of at least 80% and have at least 25 trials remaining in each experimental condition after artifact rejection (out of 40 in total)^4^. Two participants were excluded from the analyses due to low accuracy; three additional participants were excluded because of too few trials remaining after artifact rejection. The data of one participant was not analyzed because this participant turned out to be over 35 years old. The data of one further participant was lost due to a technical error. In total, 7 participants were excluded from the analyses leaving 33 valid datasets.

The participants included in the analyses were on average 22.2 years old (range 19-28); 10 were male, 23 female; 29 were right- and 4 were left-handed. We chose to include left-handed participants even though it is not conventional in neuroimaging language research because the likelihood of their language function being right-lateralized is only marginally higher than for right-handed participants (Mazoyer et al., 2014). However, to ensure that we are not observing a specific pattern of results due to inclusion of the left-handed individuals, we additionally ran all analyses in this project excluding the left-handed individuals. In none of the analyses did the results substantially change after excluding the 4 left-handed participants.

### Materials

The present study had a 4 x 2 design with the factors adjective type (scalar, intersective, pseudoadjective, letter string) and noun type (low specificity, high specificity). Examples of experimental materials are provided in *Table 1*; all materials and their properties are available for download in the *Supplemental online materials* (https://osf.io/kyc4u/). The scalar, intersective, and letter string conditions were included as a replication of the Z&P design, whereas the pseudoadjective condition was added in this study in order to investigate syntactic composition with the meaning stripped away. Note that, for easier comparison, in this section and below we provide information about the Z&P set-up in the footnotes whenever we diverged from it.

**Table 1.** Examples of materials in each experimental condition

| condition | components | adjective | low spec noun | high spec noun |
| --- | --- | --- | --- | --- |
| scalar adjective | sem + syn | klein(e) [small] | insect [insect] | vlinder [butterfly] |
| intersective adjective | sem+syn | dood(e) [dead] |  |  |
| pseudoadjective | syn | eelm(e) |  |  |
| letter string | none | gcxp |  |  |

We selected 40 noun pairs where one noun had a more general meaning - the low-specificity condition, e.g., *insect* [insect], *tas* [bag], *groente* [vegetable] - and the other noun had a more specific meaning - the high-specificity condition, e.g., *vlinder* [butterfly], *slaapzak* [sleeping bag], *wortel* [carrot]. The high-specificity noun was always in a set-theoretic subset relation to the low-specificity noun according to the categorization in WordNet (PrincetonUniversity, 2010). Low- and high-specificity nouns were matched on log10 frequency (SUBTLEX-NL; Keuleers, Brysbaert, & New, 2010)^5^, lexical decision times (Dutch lexicon project; Brysbaert, Stevens, Mandera, & Keuleers, 2016), number of letters and number of morphemes. In Dutch, nouns have a grammatical gender property - they can be either of *common* or *neuter* gender; it was not possible to match the number of low- and high-specificity nouns in grammatical gender exactly, but they were approximately matched. Summary of noun properties is provided in *Table 2* ^6^.

**Table 2.**
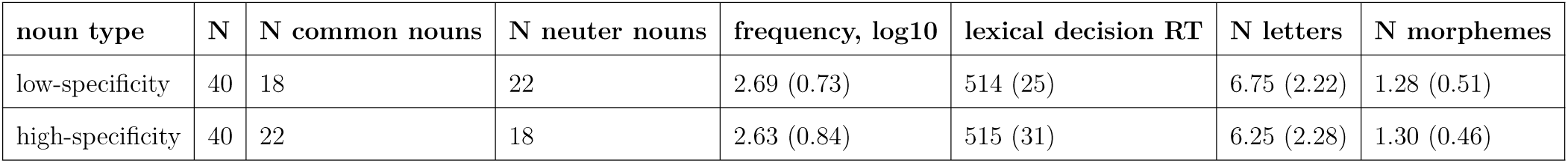
Noun properties. The value in the brackets indicates standard deviation.

Each of the nouns was presented in all 4 conditions, i.e., in combination with a scalar adjective, an intersective adjective, a pseudoadjective and a letter string. We used 20 unique intersective adjectives, each combined with 2 nouns; 19 unique scalar adjectives of which 17 were combined with 2 nouns and 2 were combined with 3 nouns; 20 unique pseudoadjectives and 20 unique letter strings also combined with 2 nouns each. The scalar and intersective adjectives themselves could not be matched on frequency and length since scalar adjectives are substantially more frequent and shorter than intersective adjectives^7^; see summary of properties in *Table 3*.

**Table 3.** Adjective properties. The value in the brackets indicates standard deviation.

| Adjective type | N (unique) | frequency, log10 | N letters, common form | N of letters, neuter form |
| --- | --- | --- | --- | --- |
| scalar | 19 | 3.66 (0.60) | 5.11 (0.66) | 4.11 (0.66) |
| intersective | 20 | 2.91 (0.78) | 8.40 (2.14) | 7.55 (2.01) |
| pseudoadjective | 20 | - | 5.20 (0.89) | 4.15 (0.67) |
| letter string | 20 | - | 5.10 (0.91) |  |

The combination of each noun with each adjective condition resulted in 320 experimental trials in the main experiment ^8^. The adjective-noun phrases were matched on a number of properties; see summary in *Table 4*. For real adjective conditions, initially we generated a list of 259 different adjective-noun combinations, which entered a plausibility pre-test. In this pre-test, we asked participants to give each phrase a score from 1 to 7 on how natural it sounds (more details about methods and participants, data collection and analysis code for each pre-test are available in the *Supplemental online materials*). Every adjective-noun combination received a score from 25 Dutch native speakers recruited in a web-based study. Based on the average scores, we selected 80 scalar adjective-noun phrases and 80 intersective adjective-noun phrases with a matched average plausibility score^9^. In the next step, we wanted to ensure that scalar and intersective adjectives we used are indeed perceived as different classes of adjectives in terms of their context-sensitivity. One possible test of scalarity that was used by Z&P is embedding an adjective in a phrase like ‘large for an insect’ where a scalar adjective forms a meaningful phrase whereas an intersective one does not (e.g., #‘dead for an insect’; Kamp & Partee, 1995; Siegel, 1976), presumably related to the fact that ‘for a’ requires retrieving a comparison class which is not possible for properties denoted by intersective adjectives (i.e., there are no different levels of being ‘dead’). To test this, we followed Z&P and collected data in a pre-test where participants saw each adjective and noun combination in this form: ‘[adj] for a [noun]’, and were asked to judge how natural they sound. We obtained scores for each phrase from 20 different Dutch native speakers in a web-based study. The intersective adjectives received a mean score 2.8, whereas scalar adjectives received a mean score 5.8.^10^ The real adjective-noun combinations were also matched on transitional probability from the adjective to the noun, i.e., probability that the adjective is followed by the target noun (calculated based on Dutch Google web n-gram corpus (Brants & Franz, 2009), using a web-interface provided by Information Science department of the University of Groningen). Specifically, transitional probability was defined as adjective-noun bigram frequency divided by the total frequency of the adjective in the corpus minus the cases where the adjective was at the end of the sentence or where it was followed by punctuation. Finally, real adjective-noun combinations were matched between conditions on cosine similarity (similarity scores were computed using R package *LSAfun* (Günther, Dudschig, & Kaup, 2015) based on vectors for each word created using a skipgramm-model trained on Dutch Wikipedia texts taking into account morphology (Grave, Bojanowski, Gupta, Joulin, & Mikolov, 2018).

**Table 4.** Adjective-noun phrase properties. The value in the brackets indicates standard deviation.

| adjective type | noun type | transitional probability | cosine similarity | plausibility | scalarity |
| --- | --- | --- | --- | --- | --- |
| scalar | low-spec | 0.0009 (0.0021) | 0.20 (0.06) | 6.02 (0.82) | 5.55 (0.74) |
|  | high-spec | 0.0005 (0.0010) | 0.22 (0.07) | 6.15 (0.52) | 6.06 (0.57) |
| intersective | low-spec | 0.0015 (0.0037) | 0.24 (0.13) | 5.88 (0.81) | 2.93 (0.76) |
|  | high-spec | 0.0011 (0.0024) | 0.24 (0.10) | 5.89 (0.81) | 2.69 (0.78) |

The pseudoadjective condition in our study was conceived as the condition where we should observe morphosyntactic composition processes but no meaning/semantic composition. The ending of the pseudoadjective always agreed with the following noun in terms of grammatical gender marking. The nonwords (pseudoadjectives) were generated based on real words in CELEX database using WordGen software (Duyck, Desmet, Verbeke, & Brysbaert, 2004), making sure that each 2-letter bigram in the nonword actually appears in a real word in Dutch at least 30 times. Afterwards, the ending ‘e’ was added to these nonwords in cases when they were combined with a noun of common grammatical gender, and no ending when combined with a noun of neuter grammatical gender. In order to discourage semantic composition, we had to be sure that participants do not have any meanings associations with the pseudoadjectives that we used. To test for that, we administered a pre-test in which participants were presented with a pseudoadjective and 4 options for nouns that could follow it. The participants’ task was to click on the noun that their intuition told them best matched as a continuation of this adjective (the pseudoadjectives were interspersed with real adjectives). One of the 4 possible options was the target noun (i.e., the noun we wanted to use in the experiment) while the other 3 options were randomly selected for every participant and every pseudoadjective from the set of all other gender-compatible target nouns (i.e., nouns that were used as targets for other pseudoadjectives). We reasoned that if a certain pseudoadjective-noun combination has a stable association for participants, they should consistently select the target noun. If there is no stable meaning association, the target noun should be as suitable as the distractor nouns and, hence, be selected at a chance level - around 25% of the time. We obtained judgments from 30 Dutch native speakers for each pair in a web-based study. Because of the low sample size^11^, to allow for a margin of error we considered all cases where the target noun was selected by 30% or less of the participants corresponding to a chance level. Since it turned out to be difficult to avoid any pseudoadjective-noun pairs that were selected by more than 30% of participants, we decided to include 4 pseudoadjective-noun pairs (out of 80 in total) which exceeded this threshold. In the selected pseudoadjective-noun pairs, the average proportion of participants who selected the target noun was 21% (range 6-37%).

Another important consideration with pseudoadjective-noun combinations was that since these were nonwords without meaning and there were no function words presented, we could not be sure that participants would indeed carry out the morphosyntactic composition with the nouns for them. In principle, they could also ignore pseudoadjectives and still respond to comprehension questions (see *Procedure* below). As a way to find out whether participants engaged in morphosyntactic composition for pseudoadjective-noun combinations, we added an additional block of 60 trials at the end of the main experiment, i.e., in addition to the main 320 trials. Out of these, in 20 trials the ending of the pseudoadjective mismatched the grammatical gender of the noun (violation condition), in 20 trials the ending of the pseudoadjective matched the grammatical gender of the noun (correct condition) and in 20 filler trials with real adjectives, the adjective-ending agreed with the grammatical gender of the noun. These trials had exactly the same structure as in the main experiment. If participants indeed engage in morphosyntactic processing for the pseudoadjective conditions, we expected to observe an event-related fields (ERF) signature of violation processing when contrasting the violation and correct grammatical gender marking conditions. If participants did not engage in morphosyntactic processing, their processing system should not be sensitive to the mismatch and, therefore, we should not observe a differing ERF pattern. There is no parallel previous study that has looked at ERF or ERP signatures in response to violations of grammatical gender marking on pseudowords but there is some research that looked at similar effects. Some studies with real words looked at gender agreement violations in adjective-noun phrases (although none in Dutch), and report more negative ERPs for violations approximately 300-500 ms after onset of the mismatching word (the so-called late anterior negativity effect) followed by more positive ERPs for violations approximately 500-800 ms after the onset of the mismatching word (the so-called P600 effect) (Molinaro, Barber, & Carreiras, 2011). Other studies looked at gender agreement violations in determiner-noun phrases in Dutch and also report similar negativity and positivity effects (Hagoort, 2003; although Hagoort & Brown, 1999 report only the positivity effect). Based on these reports, in our study we expected to observe a difference in ERFs for nouns preceded by pseudoadjectives with mismatching gender marking and for nouns preceded by pseudoadjectives with matching gender marking approximately in the 300-500 ms after noun onset time-window. We could not look at later effects since at 600 ms after noun presentation the participants already saw the comprehension question of that trial (since we used the same trial structure for this additional block of trials).

Finally, the letter string condition was intended only as a control with the same amount of visual input but no semantic or syntactic information. For this condition, we simply generated strings of consonants and ensured that they did not form phonotactically legal Dutch words.

### Procedure

During the experiment, the participants were seated in a dimly lit magnetically-shielded room, in a chair with the stimuli projected approximately 50 cm away from their eyes. They gave responses using a button-box. MEG data were acquired with a 275-channel whole-brain axial gradiometer system (CTF VSM MedTech) at a sampling rate 1200 Hz with an analog 300 Hz low-pass filter. The position of the participants’ head was recorded and monitored by the experimenter in real time throughout the recording, using 3 coils placed on the nasion, and in the left and right external auditory meatus (Stolk, Todorovic, Schoffelen, & Oostenveld, 2013); in case the participant moved their head more than 10 mm from the original position, they were asked to return to it during the next break (see below). Bipolar vertical and horizontal EOG as well as ECG were recorded using Ag/AgCl-electrodes (placed above and below the left eye, to the left of the left eye and to the right of the right eye, on the right shoulder (in supraclavicular fossa) and on the lowest left rib) with an electrode placed on the left mastoid as the ground electrode.

After the MEG recording, an MRI scan for each participant was obtained on one of three 3T MRI scanners (Siemens) available at the Donders Institute using a high-resolution T1-weighted magnetization-prepared rapid gradient-echo pulse sequence. For the MRI acquisition, the participants were wearing the same in-ear moulds as for the MEG recording, but with a Vitamin E tablet in them; these points in the ear canal along with the nasion were later manually marked and used for co-registration of anatomical data to the head location during MEG recording. The participants’ head shape was digitized using a Polhemus Fastrak digitizer, and also used for coregistration.

Stimulus presentation was controlled by software package Presentation (Neurobehavioral Systems). The experiment started with the display of instructions, and the participants had a chance to practice on 4 trials. Participants were instructed to ‘read the combinations of words and answer questions about them’. They were told that sometimes the first word would be a real word and sometimes not; when the first word would be a real word, the question would be about the meaning of the combination of the two words, whereas when the first word would not be a real word, the question would be about the second word only. The comprehension questions were not full-blown sentences but one or several words followed by a question mark, which the participants were instructed to convert into proper questions themselves (for example, a question like ‘has legs?’ was supposed to be understood as ‘does it have legs?’; we did this in order to avoid participants spending time reading and re-reading long questions and in accordance with what Z&P did in their study). All participants saw all stimuli. The 320 trials of the main experiment were divided into 5 blocks of 64 trials in each; the order of the trials was fully randomized^12^. After these trials, the participants also saw the last block of 60 additional trials some of which contained pseudoadjective violations, with the order randomized within that block. Between blocks, the participants had a chance to rest for as long as they wanted; if needed, during the break the experimenter asked the participants to return their head to the original position.

The trial structure mirrored the one used by Z&P; see *Figure 1* for an illustration. Each trial started with a fixation cross displayed for 300 ms. The adjectives and nouns were displayed for 300 ms each with a 300 ms long blank before, between and after. The adjective and noun were displayed in the center of the screen, in white 36 pt Consolas font on gray background. The comprehension questions remained on the screen until the participant gave a response by pressing one of the two buttons - ‘yes’ or ‘no’. The participants used the thumbs of the right and the left hand to respond; the mapping of the hands to the responses was identical throughout the experiment. The expected answer to 50% of the questions was “yes” and to the other 50% it was “no”. The participants were instructed to avoid blinking during the display of the words and to try to only blink during the question display.

**Figure 1.**
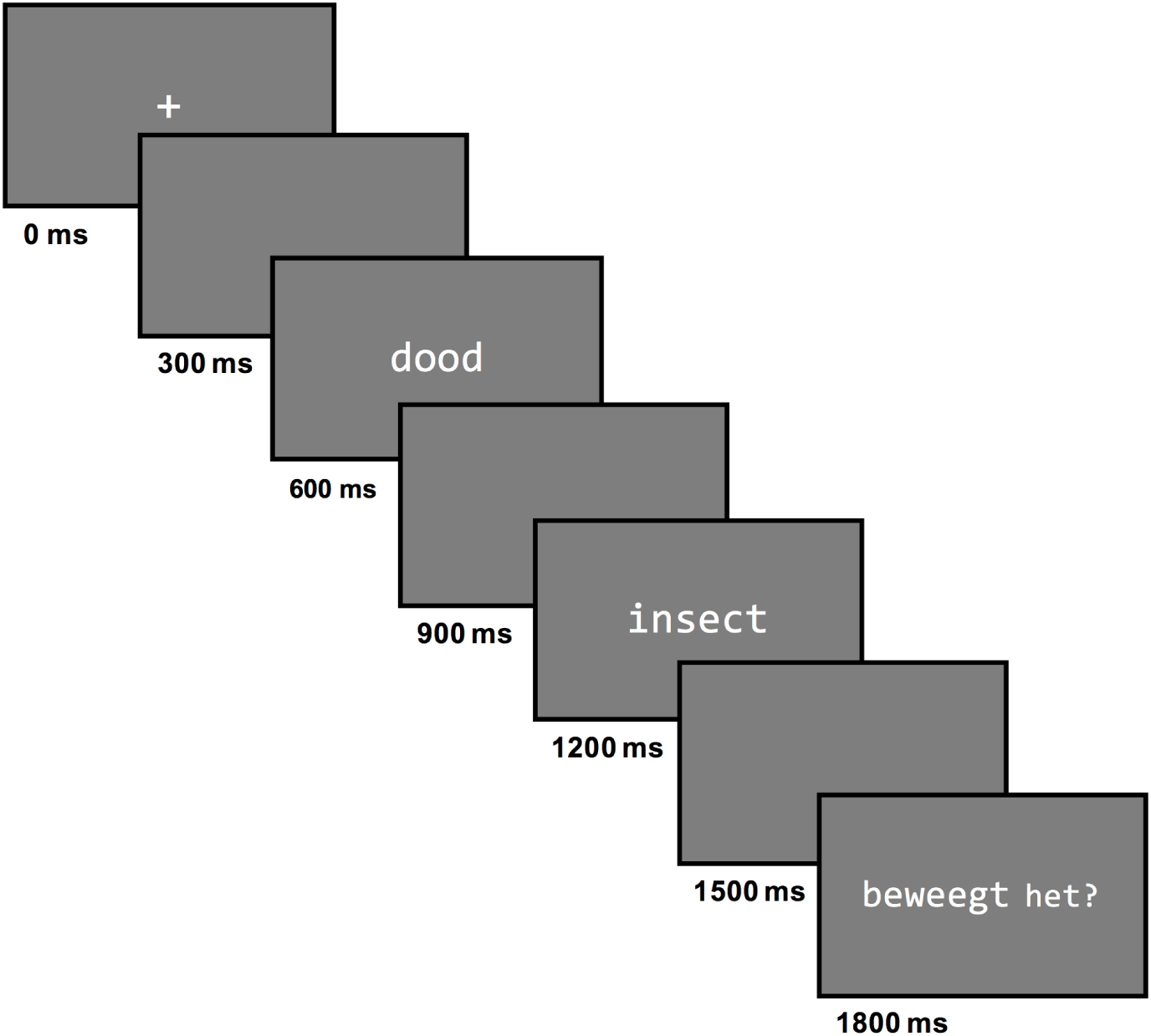
Trial structure.

The MEG recording itself took approximately 30 minutes. Each session, including preparation, practice, MEG recording and MRI scanning took approximately 1 hour and 40 minutes.

### Data pre-processing

#### MEG data pre-processing

The MEG data was processed using FieldTrip toolbox (Oostenveld, Fries, Maris, & Schoffelen, 2011) in MATLAB 2018b environment; all analysis scripts, raw and processed data are available for download in the *Supplemental online materials.* For pre-processing and source reconstruction, we followed the steps described in Z&P as closely as possible, deviating only in adding steps that we considered important to get a meaningful outcome even though they were not mentioned in the Z&P paper (again, whenever we deviated from their analysis steps, it is explicitly mentioned in the text below). The continuous MEG data was split into epochs of 700 ms before noun onset and 600 ms after noun onset (hence, containing both adjective and noun presentation windows). The epochs containing SQUID jump artifacts were excluded. A 1 Hz high-pass filter was applied to the data. All epochs which contained signal above 3000 fT were excluded. Epochs containing eye blinks were excluded using a manual and semi-automatic artifact detection procedure. Subsequently, we used ICA to determine and exclude the components corresponding to the heartbeat signal in the data. Finally, we inspected each trial in a semi-automatic fashion and excluded trials with remaining large muscle or other artifacts^13^.

#### Anatomical data pre-processing

In order to achieve better spatial localization from MEG source reconstructions, we used individual T1-weighted MRI scans of participants to create a volume conduction model of the head and a cortical sheet based source model^14^. The volume conduction model was created as a realistically shaped single shell approximation of the inside of the participants’ skull (Nolte, 2003), using FieldTrip toolbox.

As the source model, we constructed a triangulated cortical mesh using FreeSurfer’s automatic surface extraction pipeline *recon-all* (http://surfer.nmr.mgh.harvard.edu/fswiki/ReconAllTableStableV6.0). The resulting high resolution meshes were surface-registered to a common template and downsampled to a resolution of 7842 vertices per hemisphere, using HCP workbench (Marcus et al., 2011). This procedure resulted in participant-specific source models, in which individual dipole locations could be directly compared across participants.

#### Source reconstruction

For source-level activity reconstruction, data pre-processed as described above was low-pass filtered at 40 Hz, downsampled to 1000 Hz, and baseline-corrected for 100 ms before critical word onset (i.e., either before adjective onset for the analyses of adjective time-window or before noun onset for the analyses of the noun time-window)^15^. Epochs were then averaged across trials in each condition separately. The forward solution was computed using the individual source and volume conduction models and the participant-specific gradiometer positions (Nolte, 2003).

Source activity was estimated for averaged data, in each condition separately, using L2 minimum-norm estimates (Dale et al., 2000; Hämäläinen & Ilmoniemi, 1994). As a source model, we used a cortically-constrained model consisting of 15684 dipoles which were evenly distributed across the two hemispheres. At each dipole location, the 3-dimensional (i.e. orientation unconstrained) time series of activation were combined as the root mean square (RMS) across the three cardinal directions, to yield a single unsigned time course of activity. In order to be able to investigate activity in specific Brodmann Areas (BA), as was done by Z&P, the cortical source models were parcellated into 374 areas using an adjusted version of the Conte69 atlas^16^ (Glasser & Van Essen, 2011; Van Essen, Glasser, Dierker, Harwell, & Coalson, 2012), in which each of the larger Brodmann Areas was subdivided into slightly smaller parcels; the activity of points falling within each BA region was averaged to produce a single time course of activity in that region.

#### Sensor-level data

For the sensor-level analyses, the data pre-processed as described above was low-pass filtered at 40 Hz, downsampled to 1000 Hz, and baseline-corrected for 100 ms before critical word onset (i.e., either before adjective onset for the analyses of adjective time-window or before noun onset for the analyses of the noun time-window). The trials from each condition were averaged for each of the participants. We computed synthetic planar gradients from axial gradiometer data, and combined the vertical and horizontal components of the planar gradients for easier interpretation of the group results (Bastiaansen & Knösche, 2000).

### Data analysis: Semantic composition effect and its modulation by noun specificity and adjective class

To investigate the semantic composition effect, we performed confirmatory analyses that were directly based on the findings reported by Z&P, as well as exploratory analyses, part of which were based on our expectations from the literature and part of which was unconstrained. Whereas we will be able to interpret the results from the confirmatory analyses with confidence, all results observed with exploratory analyses will need to be verified in future research to be convincing.

#### Confirmatory analyses

For confirmatory analyses, we assumed that if the effect reported by Z&P is present in the parallel set-up in Dutch, we should be able to observe it in the same region of interest and time-window. For each participant we extracted the activity in left BA21 (see Figure 2a for the exact extent of this region), the ROI used in Z&P, averaging across vertices and time points of the cluster with the largest test statistic that they identified using cluster-based permutation analysis. We then conducted analyses with the extracted activity value as the dependent variable, expecting to observe a significant difference in the same direction as reported by Z&P (analyses performed using *R* (R Core Team, 2018) and the *ez* package (Lawrence, 2016).

**Figure 2.**
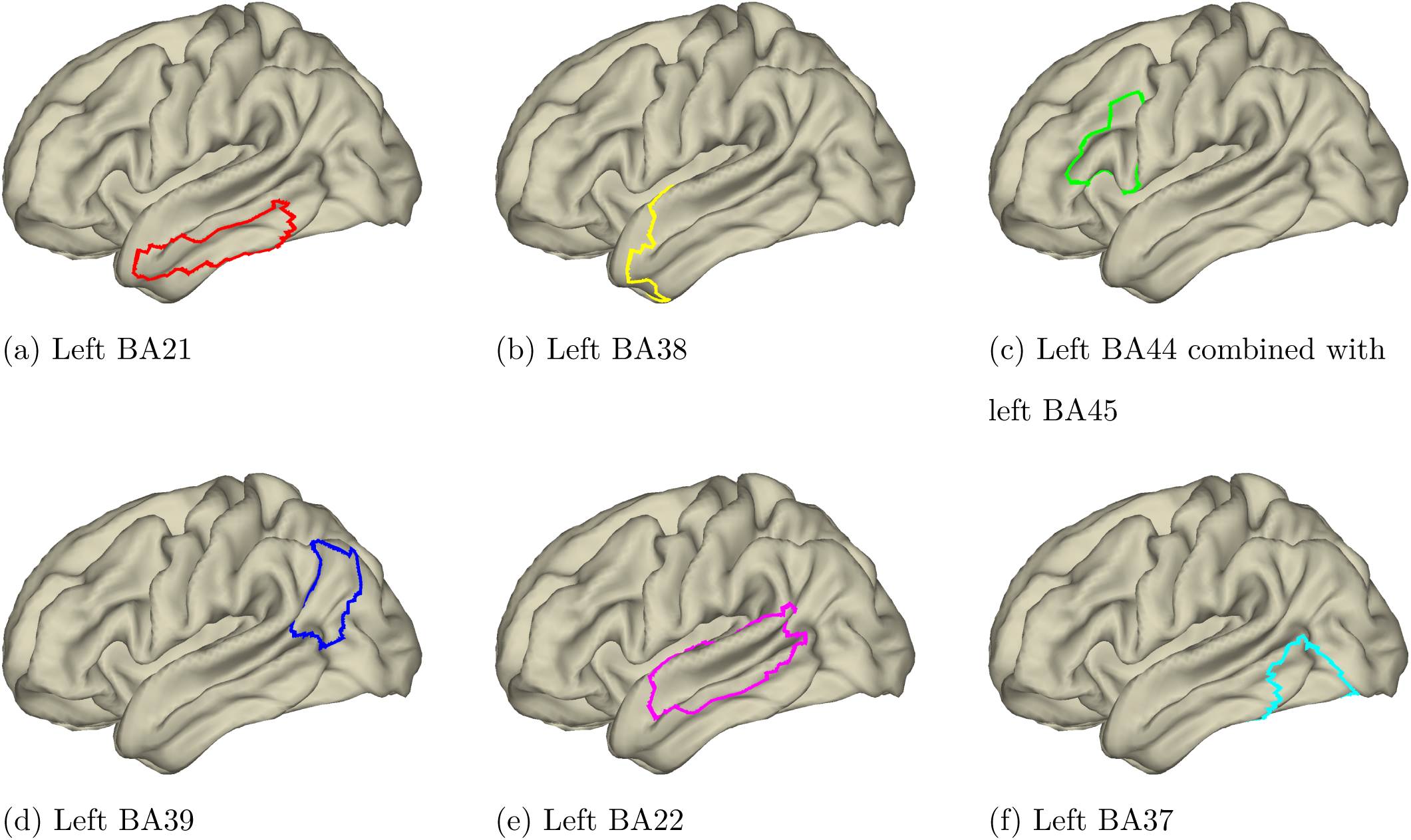
The extent of the Brodmann Areas used as regions of interest in various analyses that were conducted (based on the Conte69 atlas that was used for parcellation into BAs).

Specifically, for the LATL composition effect (i.e., difference in processing a noun preceded by a real adjective and a noun preceded by a letter string) Z&P conducted separate analyses for each adjective and noun type combination as compared to the corresponding letter string condition. They reported more activity at the noun in the condition where an intersective adjective was combined with a low specificity noun as compared to the condition where a letter string was combined with a low specificity noun; the cluster with the largest test statistic was in the time-window 200-258 ms after noun onset. They did not observe any effects when contrasting other types of adjectives and nouns with the corresponding letter string condition. We extracted the left BA21 activity values for the time-window 200-258 ms after noun onset in each of the conditions and performed 4 t-tests in parallel to their analyses, one for each of the adjective class and noun combination as opposed to the corresponding letter string and noun condition.

The second effect that we expected was a difference between adjective classes and noun types. In a 2 (scalar vs. intersective adjective) X 2 (low vs. high specificity noun) analysis, Z&P reported more activity in left BA21 for nouns preceded by intersective adjectives as opposed to nouns preceded by scalar adjectives (i.e., main effect of adjective class); with the cluster with the largest test statistic at 200-317 ms after noun onset. In addition, they reported an interaction between adjective class and noun specificity whereby noun specificity modulated left BA21 activity when nouns were preceded by scalar adjectives, but not when preceded by intersective adjectives; here the cluster with the largest test statistic was at 350-471 ms. We extracted left BA21 activity in each of these two time-windows. We ran a parallel 2×2 design repeated measures ANOVA analysis for each of the time-windows.

#### Exploratory analyses

In the confirmatory analyses, we strictly adhered to the ROI and time-window reported in Z&P study. It is, however, possible that the effect has a somewhat different timing in Dutch or that, given the variability in ROIs used in previous studies, the particular ROI that was used by Z&P does not capture the effect well. For this reason, in addition to the confirmatory analyses, we performed a series of exploratory analyses. For these exploratory analyses, we collapsed trials across different adjective classes and noun types, thus comparing simply neural activity at any noun when it was preceded by any real adjective as opposed to when it was preceded by a letter string. In case we did observe a composition effect for this contrast, we planned to investigate it further by dividing the activity level in the observed region and time-window into separate adjective and noun levels. This latter step would be used to determine whether the observed composition effect is modulated by semantic-conceptual factors, i.e., reflects semantic processing.

In the first step, we inspected a wider range of possibilities in both time and space (ROI) but still with certain constraints given what we know from the literature. Based on our review of previous studies, we chose four ROIs and performed the analysis for each of these regions separately: left BA21 that was used as an ROI in Z&P, BA38 (see Figure 2b for the exact extent of this region), a region which was analyzed as the ROIs in many of the other MEG studies (e.g., Blanco-Elorrieta & Pylkkänen, 2016; Del Prato & Pylkkänen, 2014; Pylkkänen et al., 2014; Zhang & Pylkkänen, 2015), left inferior frontal gyrus - specifically BA44 and BA45 (see Figure 2c for the exact extent of this region) - that was reported as the locus of the composition effect in an fMRI study (Schell et al., 2017), and, finally, left angular gyrus (BA39; see Figure 2d for the exact extent of this region) which was also implicated by fMRI studies (Price et al., 2015; Schell et al., 2017). For activity in each of these regions, we ran a paired t-test based cluster-based permutation analysis (Maris & Oostenveld, 2007), looking for significant differences in the time-window 100-500 ms after noun onset^17^ to capture a wider range of possible time-windows. We still expected the composition effect to manifest itself as more activity in case a real adjective is combined with a noun than in case a letter string is combined with a noun; thus, our tests were one-sided. In these analyses, consecutive time-points were grouped into clusters when they showed an effect at an uncorrected level p<0.3^18^. A corrected, Monte-Carlo significance probability was calculated as the proportion of 10 000 random permutations yielding a cluster with a higher test statistic than the cluster with the highest test statistic in the actual data. Given that we ran four separate tests, we Bonferroni corrected the significance level to be 0.05/4 = 0.0125. Note that because in these analyses we collapsed data across different levels of adjective class and noun specificity, we had an unequal number of trials in the two conditions (we had 160 trials with real adjective-noun phrases but only 80 trials with letter string-noun phrases). In order to equalize signal-to-noise ratio in the data, we performed source reconstruction for the real adjective-noun phrase condition using a randomly selected subsample of real adjective trials for each participant, with the number of trials here being the same as the number of trials available in the letter string-noun phrase condition for the same participant. To ensure that we are not missing anything because of a specific selected subsample of real adjective-noun phrase trials, we ran 100 different iterations of the analysis, each time selecting a new random subsample of real adjective-noun phrase trials for source activity reconstruction. If an effect is present in at least 80% of the analysis iterations, we would consider this effect reliable.

In the second step, we conducted an unconstrained exploratory analysis looking for potential composition effects in the whole brain at the level of individual dipole sources. Here, we conducted cluster-based permutation analysis looking for both spatial and temporal clusters in the time-window 100-500 ms after noun onset. This test was also one-sided, looking for more activity in the condition where a real adjective is combined with a noun in comparison to a condition where a letter string is combined with a noun. Cortical locations and time-points that showed an effect at an uncorrected level p<0.05 were grouped into clusters. A corrected p-value was calculated based on 1000 permutations (the number of permutations was lower than in other analyses presented here since this analysis is significantly more time-consuming than the spatially constrained ones). We considered p-values below 0.05 as significant.

### Data analysis: Syntactic composition effects

To investigate brain activity related to syntactic composition, we again performed an analysis constrained by what we know from literature, as well as a whole brain analysis. Prior to exploring the syntactic composition effects, we analysed the ERFs at sensor level at the nouns in the last block of the experiment where pseuadoadjectives with matching and mismatching grammatical gender marking were presented. If we observe an agreement violation effect when comparing matching and mismatching trials, we would be convinced that our participants performed syntactic composition in the pseudoadjective condition. In absence of such violation effects, we cannot be sure that syntactic composition indeed happened in this condition as we intended, so any effects that we observe in the analysis intended to look at syntactic composition cannot be confidently interpreted as reflecting necessarily syntactic composition.

#### Morphosyntactic agreement violation

The ERFs were calculated at the noun presentation time-window of trials in the last block, for the condition where the noun was preceded by a pseudoadjective with a matching grammatical gender and for the condition where the noun was preceded by a pseudoadjective with a mismatching grammatical gender. We performed a paired t-test based cluster-based permutation analysis comparing ERFs, looking for spatiotemporal clusters between 100-600 ms after noun onset. For cluster selection, time-points and sensors showing an effect at an uncorrected level p<0.05 were grouped into clusters; minimum two neighbouring channels showing an effect were required to form a cluster. A corrected significance probability was calculated based on 5000 random permutations.

#### Exploratory analyses

We compared source-reconstructed neural activity for processing a noun preceded by a pseudoadjective as opposed to when preceded by a letter string. Based on our literature review, we identified three candidate regions in which we expected to observe a syntactic composition effect: left inferior frontal gyrus (BA44, BA45), left anterior/middle temporal lobe (BA21) and left posterior temporal lobe (BA22; see Figure 2e for the exact extent of this region). For each of these regions separately, we ran an analysis with the same set-up as the one described for semantic composition effects above, expecting higher activity levels for the nouns preceded by pseudoadjectives as compared to the nouns preceded by letter strings. Because the number of trials in each condition was approximately equal, there was no need to subsample trials and run multiple iterations of this analysis. We looked for temporal clusters between 100 and 600 ms after noun onset. Given that we conducted a separate test for each of the regions, the significance criterion level was Bonferroni corrected to 0.05/3 = 0.016.

We subsequently ran an analysis looking for potential syntactic composition effect in the whole brain, at the level of individual dipole sources. We expected to observe more activity for nouns preceded by pseudoadjectives as compared to nouns preceded by letter strings. We conducted analysis looking for spatial and temporal clusters in the time-window 100-600 ms after noun onset, with the same set-up as for the whole brain analysis for the semantic composition effect.

## Results

### Behavioral results

Participants included in the analysis gave an expected response to the comprehension questions on average on 91% of trials (SD 3.4; range 83%-96%). The mean response time was 1.56 seconds (SD 0.56; range 0.8-3.3 seconds). The purpose of the comprehension questions was only to make sure that participants pay attention to the task, and, therefore, no further analyses on behavioral data were conducted.

### Trial exclusion

The artifact rejection steps resulted in exclusion of 10% of trials overall (range 2-24%)^19^.

### Semantic composition effect and its modulation by noun specificity and adjective class

#### Confirmatory analyses

For the LATL composition effect with different adjective class and noun specificity combinations at the time-window 200-258 ms after noun onset, we did not observe a significant difference in any of the comparisons: intersective adjectives vs. letter strings combined with low specificity nouns (t[32]=-1.05, p=0.3), intersective adjectives vs. letter strings combined with high specificity nouns (t[32]=0.07, p=0.9), scalar adjectives vs. letter strings combined with low specificity nouns (t[32]=1.07, p=0.3), scalar adjectives vs. letter string combined with a high specificity nouns (t[32]=0.9, p=0.4). The plots depicting the levels of activity in left BA21 for each of these comparisons are provided in Figure 3. We thus did not find evidence for an LATL composition effect or its modulation by adjective class and noun specificity in the region and time-window where such an effect was reported by Z&P.

**Figure 3.**
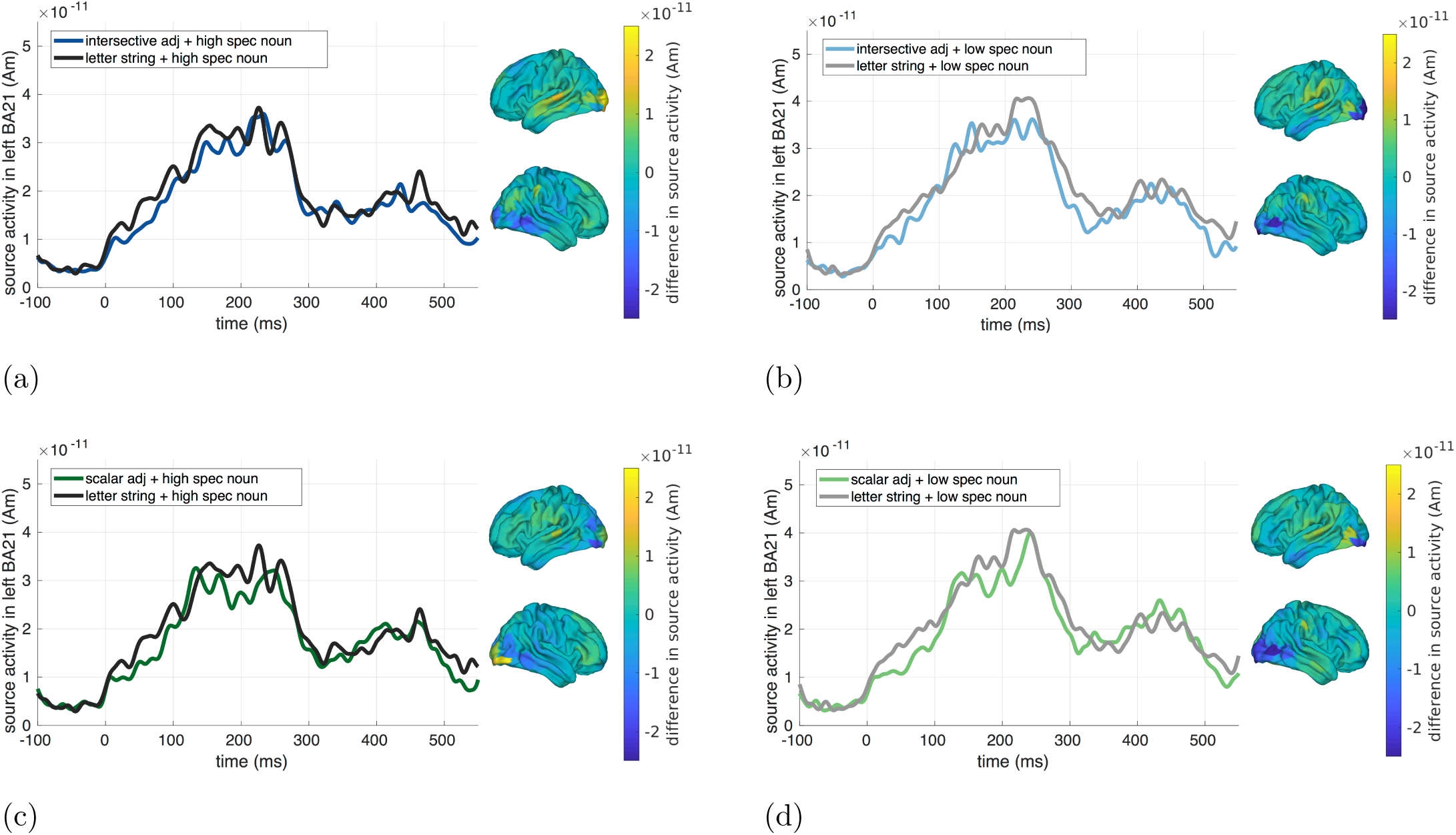
Source reconstructed activity levels in the left BA21 for conditions where the noun was preceded by a real adjective and by a letter string. Note that the time-point 0 here corresponds to the onset of the noun. The whole brain plots show activation in the corresponding real adjective condition minus activation in the letter string condition averaged between 200-250 ms after noun onset. Panel (e) displays the extent of the left BA21.

We did not observe any differences between adjectives classes and noun specificities in the time-window 200-317 ms after noun onset where we expected the main effect of adjective class (main effect of adjective class: F[1,32]=0.2, p=0.6; main effect of noun specificity: F[1,32]=3.1, p=0.08; interaction: F[1,32]=1.2, p=0.2) or in the time-window 350-471 ms after noun onset where we expected to see an interaction effect (main effect of adjective class: F[1,32]=2.9, p=0.09; main effect of noun specificity: F[1,32]=2.1, p=0.1; interaction: F[1,32]=0.2, p=0.6). Figure 4 depicts source-reconstructed activity for different levels of adjective class and noun specificity. We thus did not find evidence for differences between adjective classes and noun specificity in the region and time-windows where such effects were reported by Z&P.

**Figure 4.**
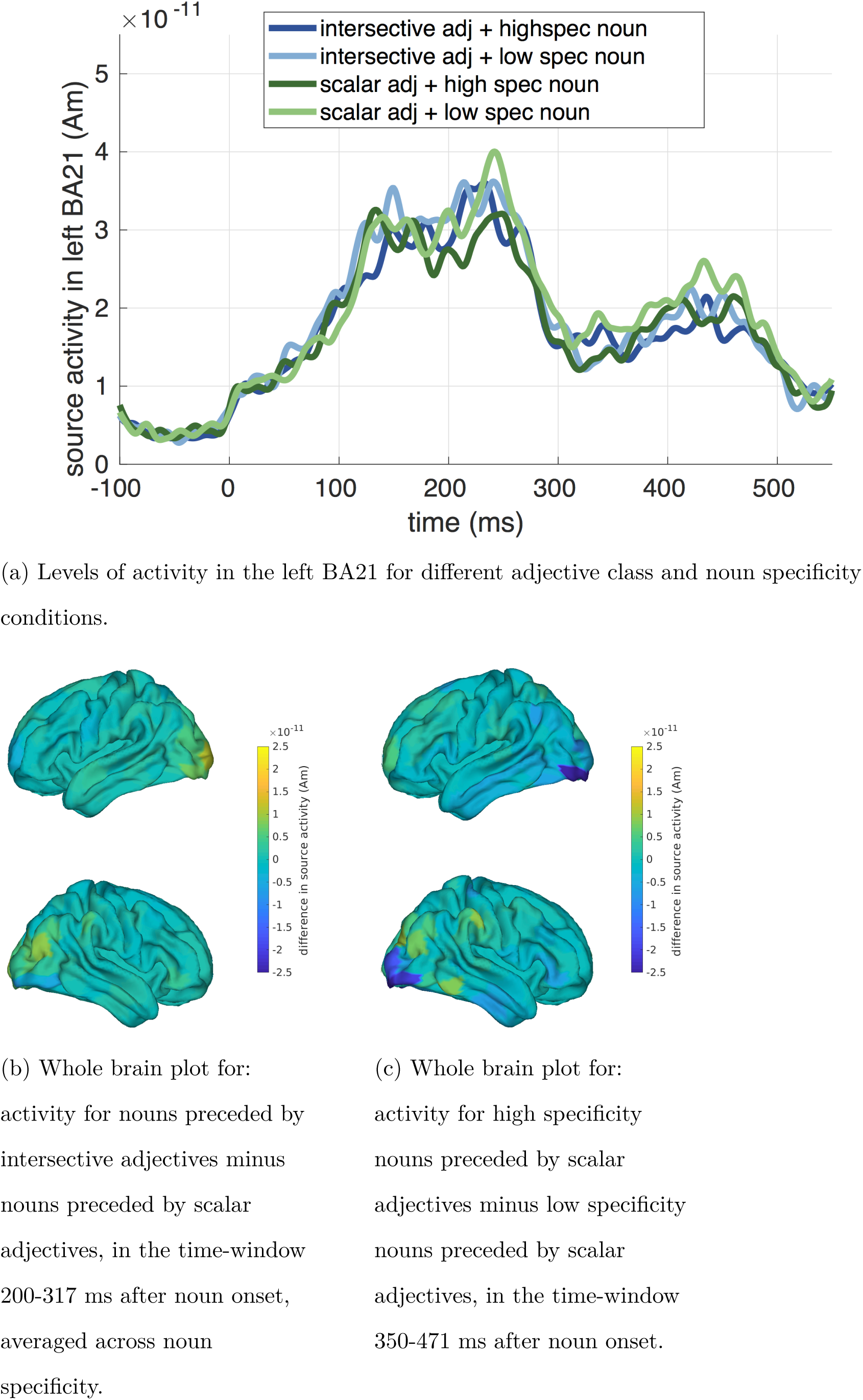
Source reconstructed activity levels for different adjective class and noun specificity conditions. Note that the time-point 0 here corresponds to the onset of the noun.

#### Exploratory analyses

In the constrained exploratory analysis, we did not observe any difference in the source reconstructed activity between a noun preceded by a real adjective and a noun preceded by letter strings in any of the regions of interest that we selected, in none of the iterations with different subsamples of real adjective trials (100 iterations). Neither did we observe any significant differences in the unconstrained exploratory analysis over the whole brain. Figure 5 depicts the activity in the whole brain for consecutive time-windows within the window of interest.

**Figure 5.**
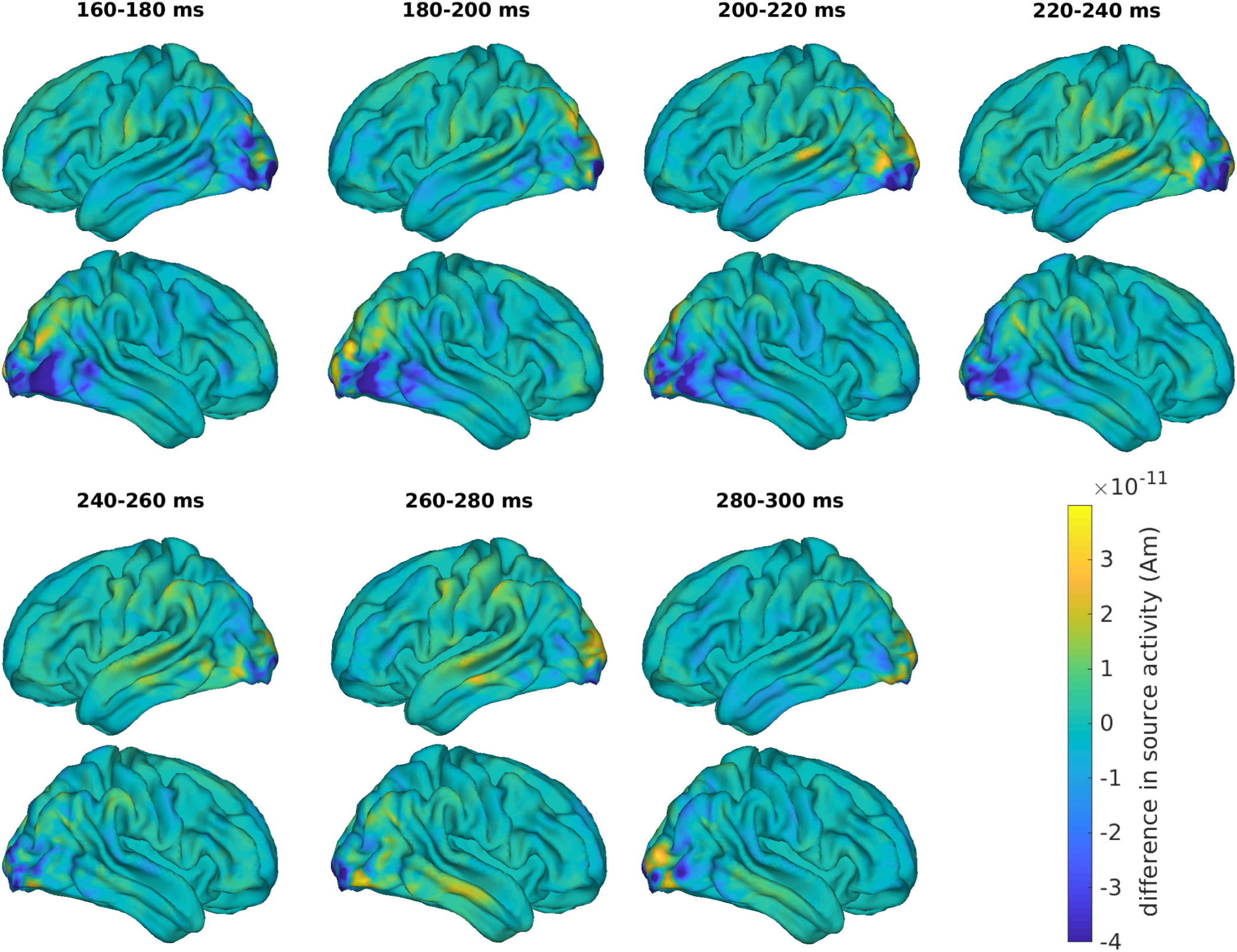
Source-reconstructed activity for nouns preceded by a real adjective minus nouns preceded by a letter string. Note that the time-point 0 here corresponds to the onset of the noun. Note that for this plot we used source-reconstructed activity from only one iteration of the analysis (i.e., one subsample of trials of the intersective and scalar adjective conditions.)

### Syntactic composition effect

#### Morphosyntactic agreement violation

We did not observe a significant difference between ERFs at the nouns preceded by a pseudoadjective with matching grammatical gender ending and those at the nouns preceded by a pseudoadjective with mismatching grammatical gender. The ERFs for each of the conditions are depicted in Figure 6. Given this result, there is no clear indication that participants noticed a mismatching agreement for the pseudoadjective-noun pairs, and we, therefore, cannot be sure that our participants performed syntactic composition in case of the pseudoadjective condition in our main experimental trials as intended. In contrast, note that we are certain that participants carried out adjective-noun composition in the conditions with the real adjectives given their behavioral performance with the comprehension questions.

**Figure 6.**
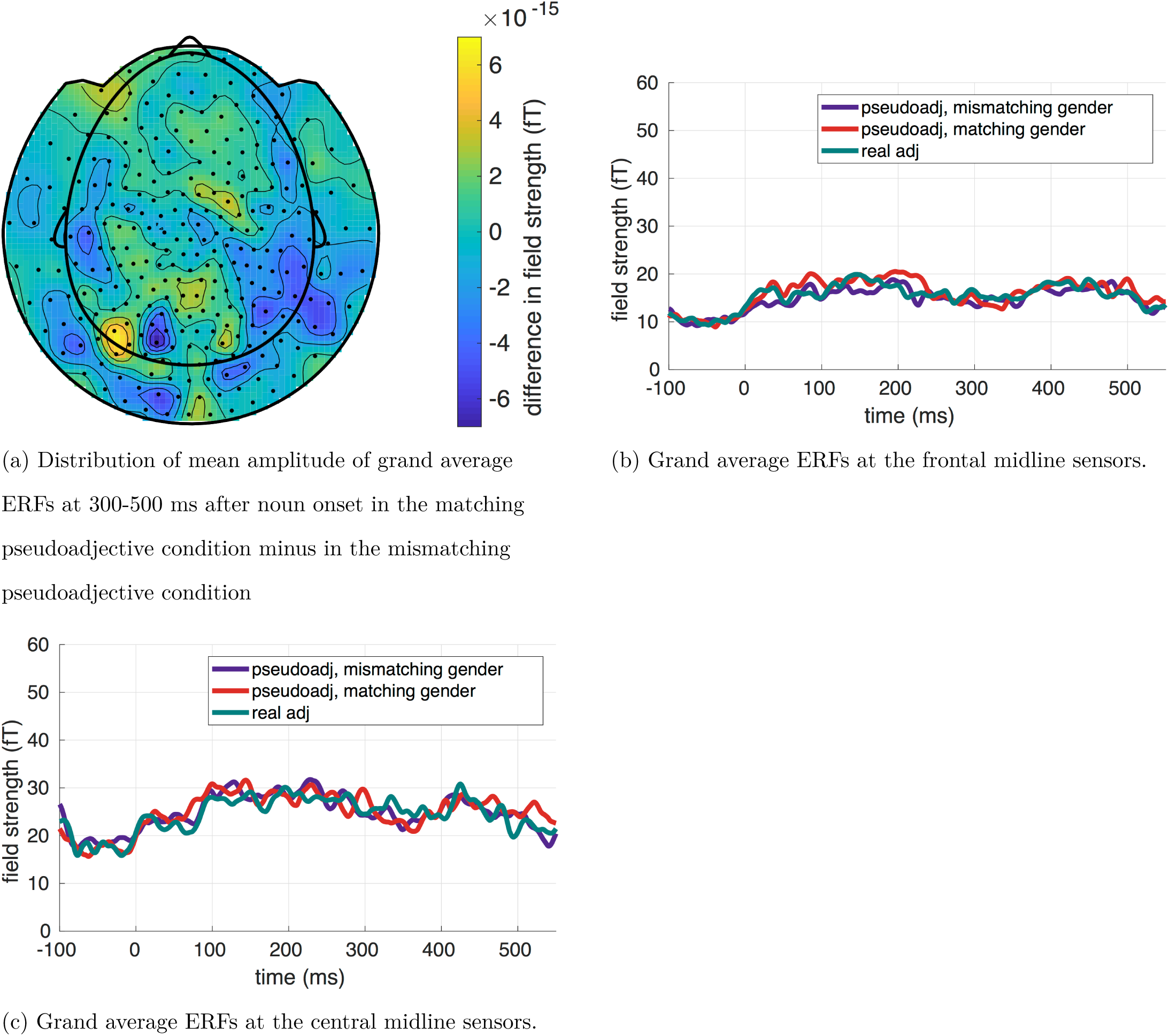
ERFs at the nouns preceded by matching pseudoadjective, mismatching pseudoadjective and real adjective (always matching) trials presented in the last, additional block of the experiment. Note that ERFs in the real adjective condition are depicted in plots B and C, but were not included in the analysis. Note that the time-point 0 here corresponds to the onset of the noun.

#### Exploratory analyses

We did not observe any difference in the source reconstructed activity between a noun preceded by a pseudoadjective and a noun preceded by letter strings in any of the regions of interest that we selected. Plots depicting activity levels in each of our regions of interest are provided in Figure 7. Neither did we observe any significant differences in the unconstrained exploratory analysis over the whole brain. Figure 8 depicts the activity in the whole brain for consecutive time-windows within the window of interest.

**Figure 7.**
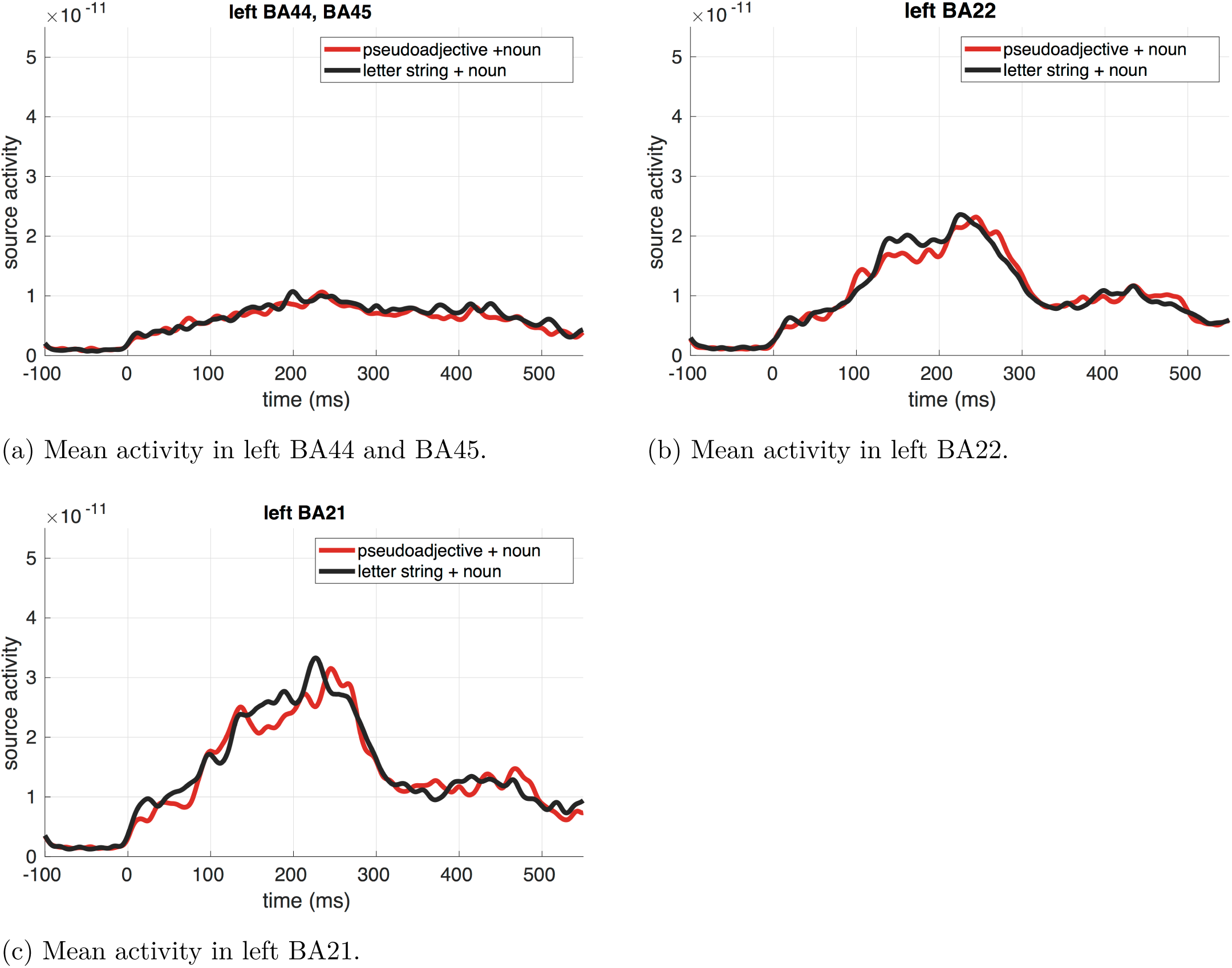
Source-reconstructed activity levels in the pseudoadjective and letter string conditions, in the regions of interest chosen for analyses of syntactic composition effects. Note that the time-point 0 here corresponds to the onset of the noun.

**Figure 8.**
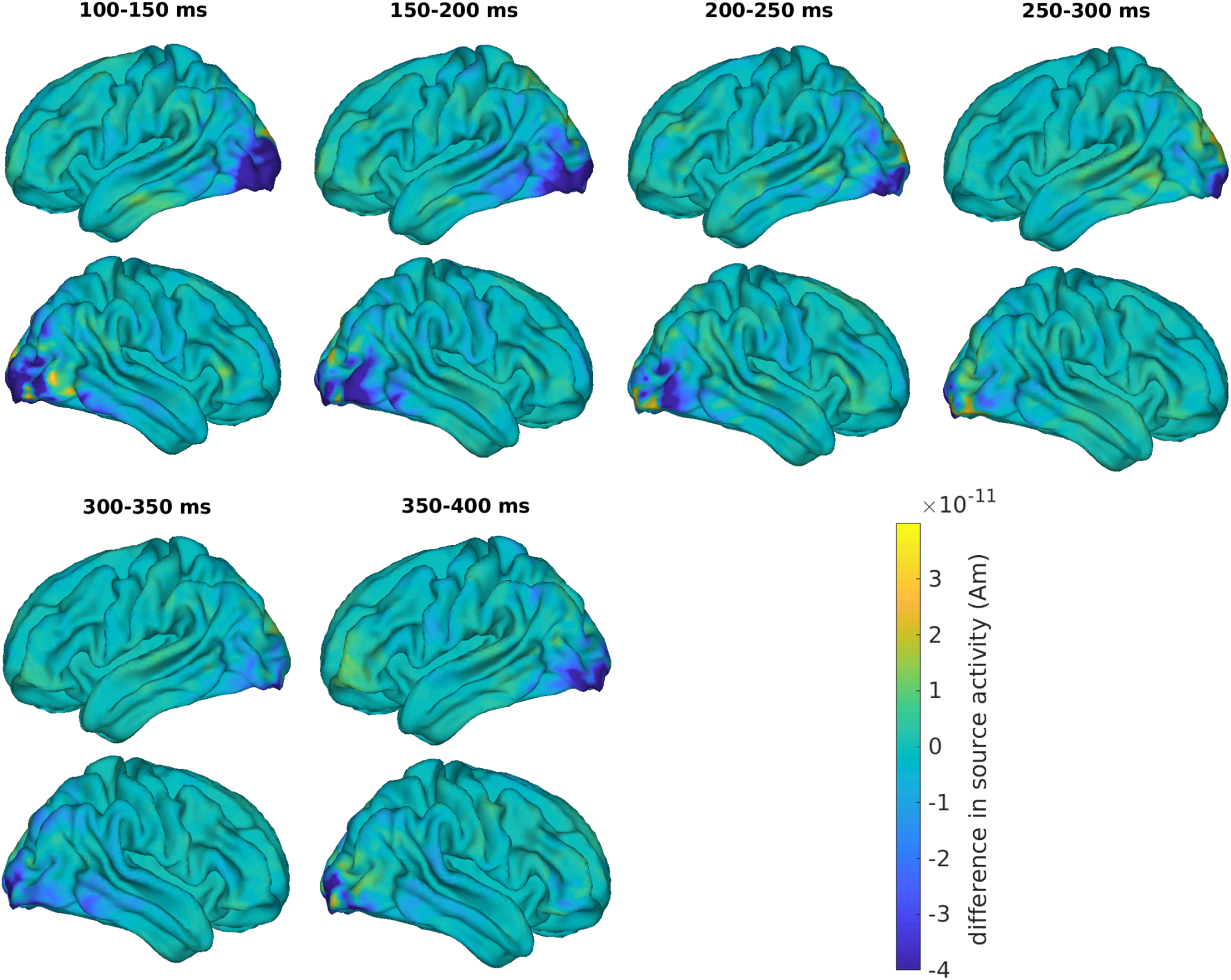
Source-reconstructed activity for nouns preceded by pseudoadjectives minus nouns preceded by letter strings. Note that the time-point 0 here corresponds to the onset of the noun.

### Validation of data processing pipeline

Before discussing our results and their implications, we take an additional step to check the validity and soundness of our data processing pipeline and analysis procedures. To do so, we looked for independent effects in our data, based on a comparison between real adjectives and pseudoadjectives. We call these ‘sanity check’ analyses.

We compared the brain response to the presentation of real adjectives and the response to the presentation of pseudoadjectives or letter strings, at a time-point before the participants saw the noun. At this point, the participants have only seen one word, so no composition effects are expected. Given their task, what participants did at this point is comparable to completing a lexical decision task: they attempted to retrieve lexical information about the word from memory, i.e., recognize the presented word. Hence, we believe that at this point we should observe effects similar to the ones that are reported for the lexical decision task where typically participants respond ‘yes’ or ‘no’ to whether a presented word is a real word.

#### For source reconstruction procedure

An MEG study by Hauk and colleagues (Hauk, Coutout, Holden, & Chen, 2012) employed a contrast that comes close to ours, and is similar in terms of the source reconstruction procedure. In their study (Experiment 2), participants were deciding whether a presented word was a real word and gave responses using eye blinks. Important differences from our study were the following: this was a pure lexical decision task (i.e., nothing followed after participants decided whether the presented word was real), stimuli were presented for just 100 ms, and the real words that participants saw were nouns. For the source-reconstructed data, Hauk and colleagues report significantly higher neural activity when processing real words as opposed to pseudowords in the left middle and inferior temporal lobes between 180-220 ms after word onset.

As a parallel contrast to this one, in our sanity check analysis we compared processing of scalar adjectives and pseudoadjectives at the time-window just following adjective onset. We did not look at intersective adjectives in this analysis because they were on average longer than pseudoadjectives (8.4 letters as opposed to length 5.1 and 5.2 letters in case of scalar adjectives and pseudoadjectives respectively) since we know that word length can influence early word processing (e.g., Assadollahi & Pulvermüller, 2003; Hauk, Davis, & Pulvermüller, 2008; Hauk & Pulvermüller, 2004; Schurz et al., 2010). Because we performed the analyses at Brodmann Area level parcels, based on visual inspection of the effect observed by Hauk and colleagues (Hauk et al., 2012, Figure 6) we estimated that we should see different activity for the two conditions in the left BA37 (see Figure 2f for the exact extent of this region) and BA22 in our parcellation. We ran a paired t-test based cluster-based permutation analysis in each of these regions, looking for temporal clusters between 100 and 300 ms. Note that the data for this analysis was baseline-corrected using a window of 100 ms before adjective onset. We ran one-tailed tests because we specifically expected to see more activity in BA37 and BA22 for scalar adjectives. The set-up of this analysis was parallel to the one used for semantic and syntactic composition effect analyses at the source-level. Given that we ran the test separately on two regions, we Bonferroni corrected the significance level to be 0.05/2 = 0.025.

As expected, we observed a higher level of activity for scalar adjectives than for pseudoadjectives in left BA37 (p=0.017) where the cluster with the largest test statistic was between 224-281 ms. For the left BA22, significance probability was 0.034, i.e., above our significance criterion; here, the cluster with the largest test statistic was between 220-258 ms. The difference in the source-reconstructed activity in the adjective presentation window for these regions is plotted in Figure 9. Note that our largest temporal cluster is at a later point than in the study by Hauk and colleagues (180-220 ms). This could be due to a different task in our study, due to a different language, or due to the fact that our materials consisted of adjectives instead of nouns. Despite the difference in the temporal extent, given that we do observe an effect with the expected spatial extent and in the expected direction, this result demonstrates that our source reconstruction pipeline produces meaningful results, reflecting activity related to language processing.

**Figure 9.**
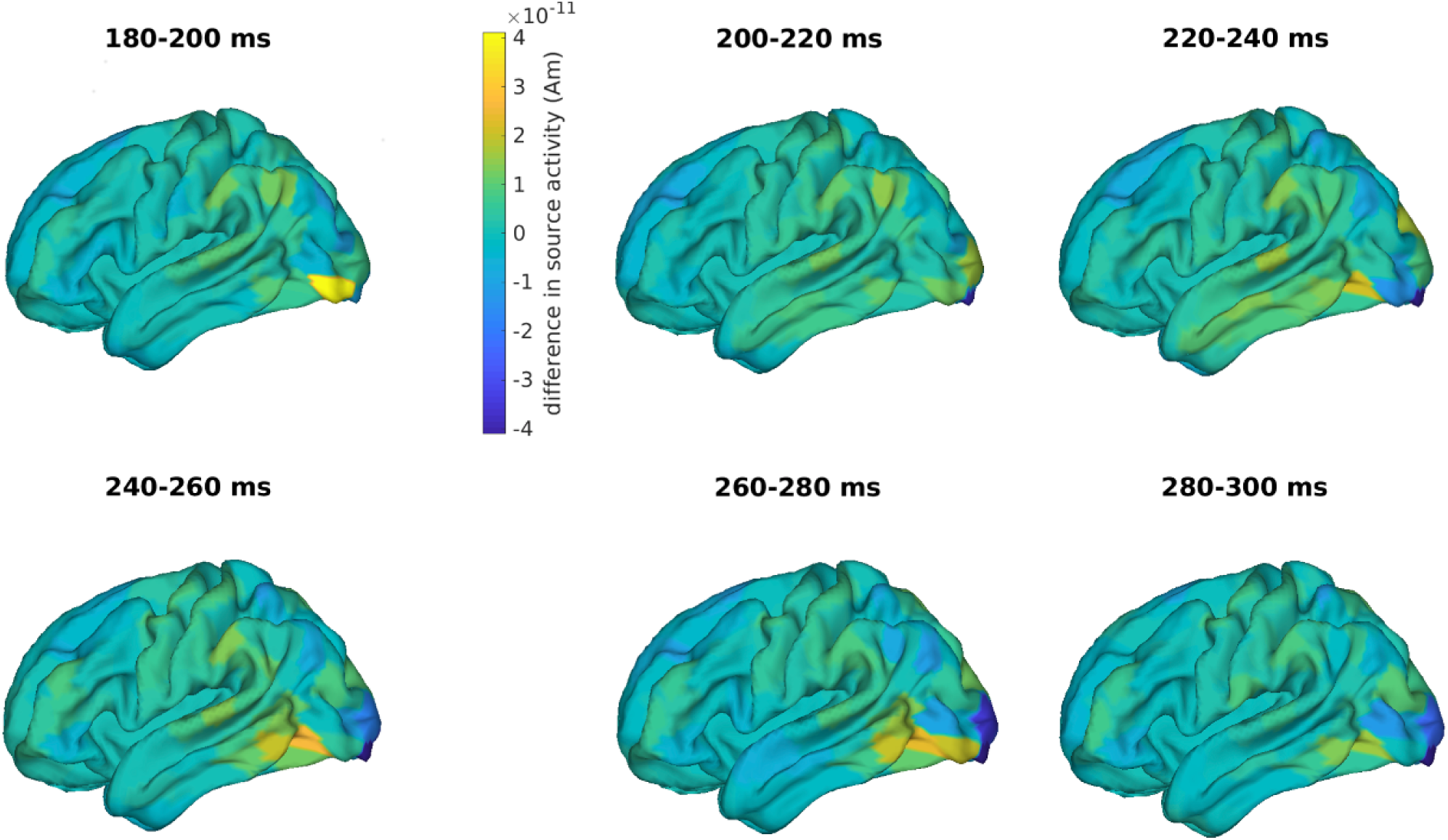
Source-reconstructed neural activity for scalar adjectives minus pseudoadjectives. Note that the time-point 0 here corresponds to the onset of the adjective.

#### For sensor-level data

We ran a second sanity check analysis at the sensor-level, as a way to validate the sensor-level ERF analysis and, thereby, provide evidence that we can meaningfully interpret the absence of a violation effect as an indication that participants might not have performed syntactic composition in the case of pseudoadjectives. One of the robust findings in the ERP literature is an N400 (or an N400-like) effect in the lexical decision task whereby a more negative ERP is typically observed for pseudowords as opposed to real words (e.g., Barber, Otten, Kousta, & Vigliocco, 2013; Bentin, McCarthy, & Wood, 1985; Kounios & Holcomb, 1994; Meade, Grainger, & Holcomb, 2019). Consistent with this, an EEG study of minimal phrase composition which employed the same paradigm and experiment structure as the original Pylkkänen lab studies and the present study (Neufeld et al., 2016) also reported such a negativity for pseudowords, at the adjective presentation window. Given those results, we thus also expected to observe an N400-like effect for pseudoadjectives contrasted with real adjectives. We again only looked at the scalar adjectives in order to equalize word length between conditions. Note that the ERF data for this analysis was baseline-corrected using a window of 100 ms before adjective onset. We ran a paired t-test based cluster-based permutation analysis comparing ERFs, looking for spatiotemporal clusters between 100-600 ms after adjective onset. The analysis was two-sided. We used the whole time-window of adjective presentation (except for the first 100 ms) in this analysis because the N400 effects for such contrasts does not have one consistently observed time-course and is often long-lasting (Barber et al., 2013; Meade et al., 2019). The set-up of this analysis was same as for the sensor-level analysis of morphosyntactic agreement violation trials.

The cluster-based permutation test revealed a significant difference between the scalar adjectives and pseudoadjectives (p=0.004). The cluster with the highest test statistic was between 368-465 ms and most pronounced over left temporal sensors; see Figure 10a. In this time-window, signal strength was greater for pseudoadjectives than scalar adjectives (see Figure 10b for depiction of grand average ERFs (note that ERFs for all conditions are depicted in the plot whereas only two were analyzed). The time and spatial features of the effect are in line with what we expected based on the results of previous studies. We can thus be confident that the pre-processing pipeline of our ERF data reflects processing of the stimuli.

**Figure 10.**
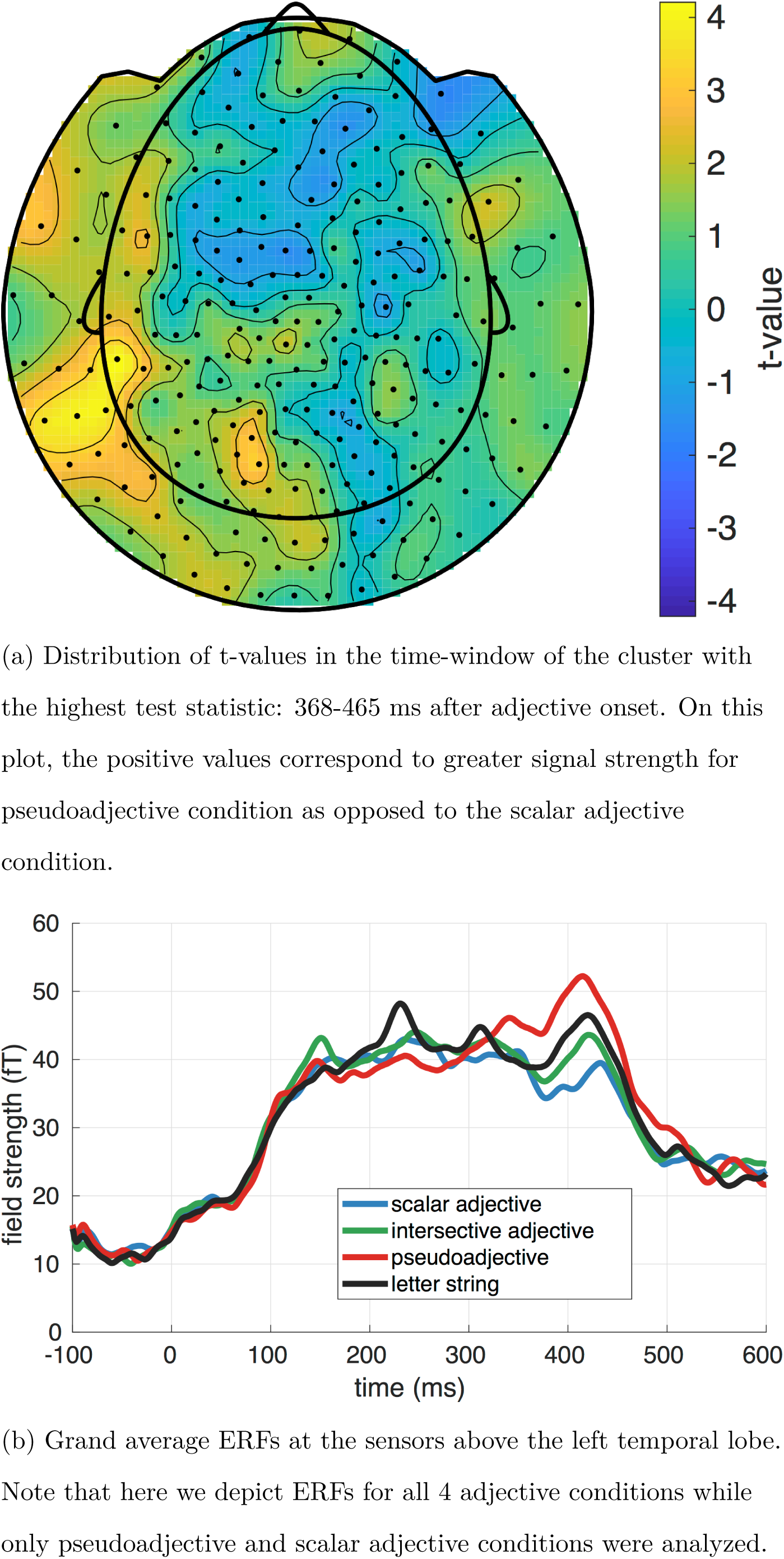
ERFs at the adjective presentation window. Note that the time-point 0 here corresponds to the onset of the adjective.

### Further exploratory analyses

In our analyses of composition effects so far, we focused on source reconstructed data following the previous MEG studies. Given that we see no differences between conditions at the source level, we investigated our data further for presence of any differences between conditions in the sensor-level data. The data at the sensor-level is noisier in comparison to source reconstructed data since multiple sources are likely contributing the signal observed at each sensor; in addition, the exact position of the participants’ heads relative to the MEG sensors is not taken into account. On the other hand, doing the analyses at the sensor-level we make fewer modeling assumptions (i.e., no need to solve the inverse problem). To our knowledge, only three previous MEG studies of LATL composition effect have reported looking at the potential differences between conditions at the sensor-level, and none of them found a significant difference (Bemis & Pylkkänen, 2011; Del Prato & Pylkkänen, 2014; Zhang & Pylkkänen, 2015). Two of these studies found a difference when narrowing down the time-window for the cluster search to just 100 ms around the point where they already report a difference in the source-reconstructed neural activity (Bemis & Pylkkänen, 2011; Zhang & Pylkkänen, 2015). Nonetheless, an EEG study looking at the composition of adjective-noun phrases with the same set-up and contrast did report a difference (Neufeld et al., 2016) so it is possible that we do see some difference as well that will help us interpret our results.

We performed sensor-level analyses in a procedure parallel to the one described for the morphosyntactic control trials and sanity check analysis. For the composition effect, we looked for a difference in ERFs between a noun preceded by a real adjective as opposed to a noun preceded by a letter string between 100 and 600 ms after noun onset. Again, to equalize signal-to-noise ratio across conditions, we performed 100 iterations of this analysis with a different randomly selected subset of real adjective trials each time. Based on the cluster-based permutation analyses, there was a difference between conditions with p-value smaller than 0.05 in 85 out of 100 iterations. Because each iteration included a different subset of trials, the temporal and spatial extent of the cluster with the largest test-statistic was slightly different in each iteration. To get the most stable time-window of the cluster with the largest test statistic, we extracted the time-window that belonged to the cluster with the largest test-statistic in at least 80% of the iterations; it was 136-180 ms after noun onset. In this time-window, the difference was most pronounced on the right central, parietal and occipital sensors; on these sensors the signal strength was smaller for the real adjective condition than for the letter string condition. The distribution of summed t-values across sensors across all 100 iterations is plotted in Figure 11a. The ERFs for the right parietal sensors is depicted in Figure 11b.

**Figure 11.**
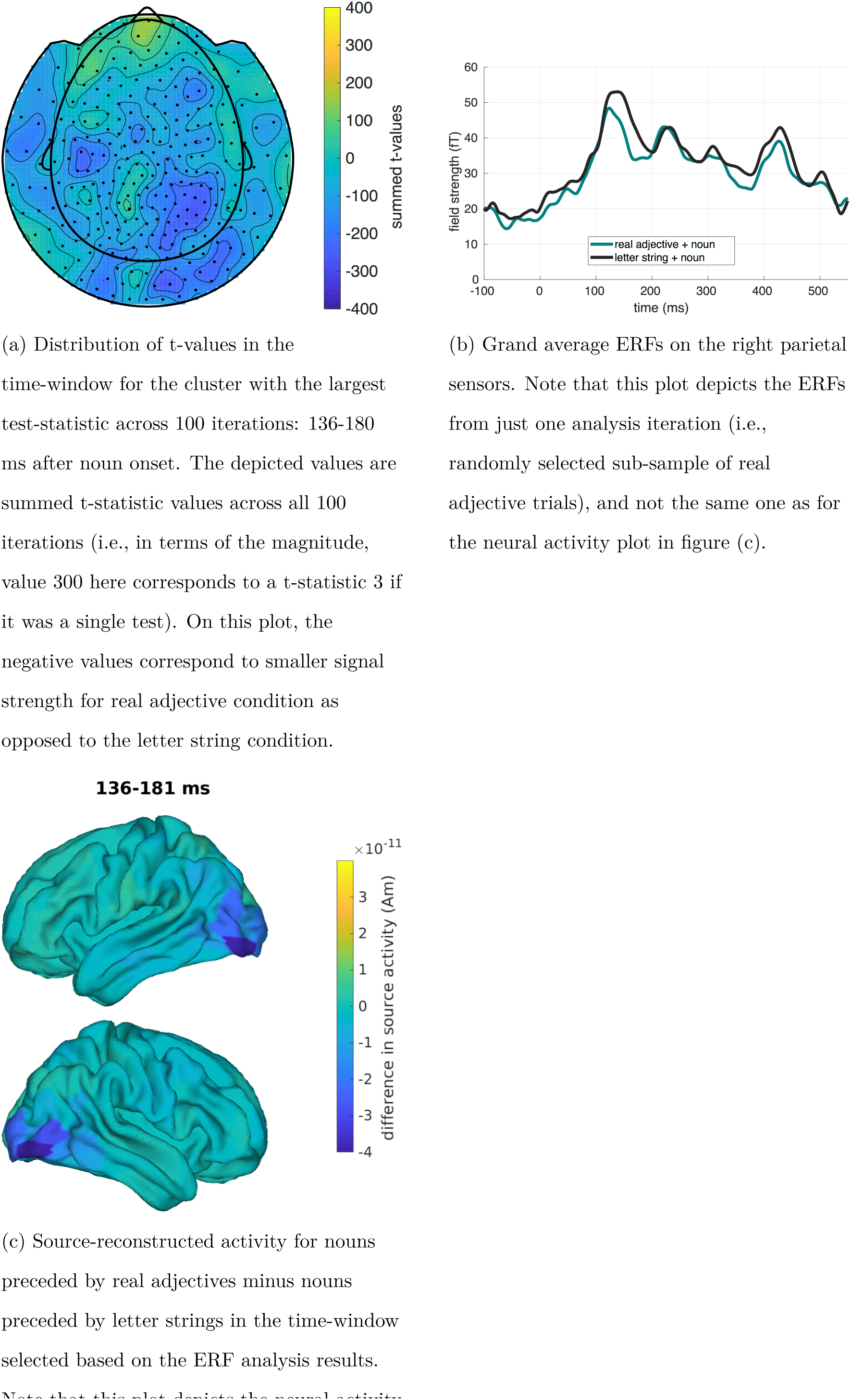
ERFs at the noun presentation window. Note that the time-point 0 here corresponds to the onset of the noun.

The time-window of this difference between conditions is earlier than that reported for the LATL composition effect (between approximately 200-250 ms after noun onset for these types of materials), so it is not clear what exactly this difference in ERFs reflects. To look at potential neural sources where the difference is originating from, we plotted and inspected the source-reconstructed neural activity in the time-window of the cluster with the largest test statistic. Figure 11c depicts the neural activity for the whole brain in this time-window. In this time-window, there was less neural activity for real adjective trials than for letter string trials in the right and left occipital lobes. This is difficult to interpret in terms of the composition effect because we expected more neural activity for the condition which requires composition processes in comparison to the condition without composition processes. In addition, occipital areas are not considered to be strongly involved for linguistic processing, but rather responsible for low-level visual processing. Given all these reasons, we believe this difference is unlikely to be reflecting composition-related processing. Rather, it is more likely that this effect reflects differences in visual processing of the noun based on what preceded it, a real adjective or a letter string.

We further looked for a potential difference in ERFs between different adjective class conditions: difference between nouns preceded by an intersective adjective as opposed to nouns preceded by scalar adjectives between 100 and 600 ms after noun onset. Note that the number of trials for each condition for this analysis was equal, so there was no need to run multiple iterations with different sub-samples of trials. The cluster-based permutation analysis did not reveal any differences between conditions.

Finally, we also conducted sensor-level analyses for the syntactic composition contrast: difference between ERFs for nouns preceded by pseudoadjectives and nouns preceded by letter strings between 100 and 600 ms after noun onset. Again, the number of trials for each condition for this analysis was equal, so there was no need to run multiple iterations with different sub-samples of trials. There was a difference between conditions with a p-value of 0.02. The cluster with the largest test-statistic was in the time-window 342-424 ms, and included the central and parietal sensors in both hemispheres. In this time-window, signal strength was smaller for pseudoadjective condition compared to the letter string condition. The distribution of t-values across sensors in this time-window is depicted in Figure 12a. The ERFs for the right parietal sensors are depicted in Figure 12b. We also looked at the potential neural sources of this effect; they too seem to be located in occipital areas. The difference in source-reconstructed neural activity between these conditions in the time-window of the cluster with the largest test-statistic is depicted in Figure 12c. This time-window matches a time-window where a linguistic N400-like effect would appear. This effect is, however, rather weak (compare it to the effect observed in the adjective time-window in the ‘sanity check’ analysis). In addition, we observe less neural activity for the pseudoadjective condition where we, in fact, expected additional processing to occur, making it difficult to interpret this effect as reflecting the syntactic composition process. Finally, the localization of this effect is, again, inconsistent with such an interpretation - occipital areas are typically considered to be responsible for low-level visual processing, and it is unclear why they would be involved here. This difference, thus, also rather reflects some other process.

**Figure 12.**
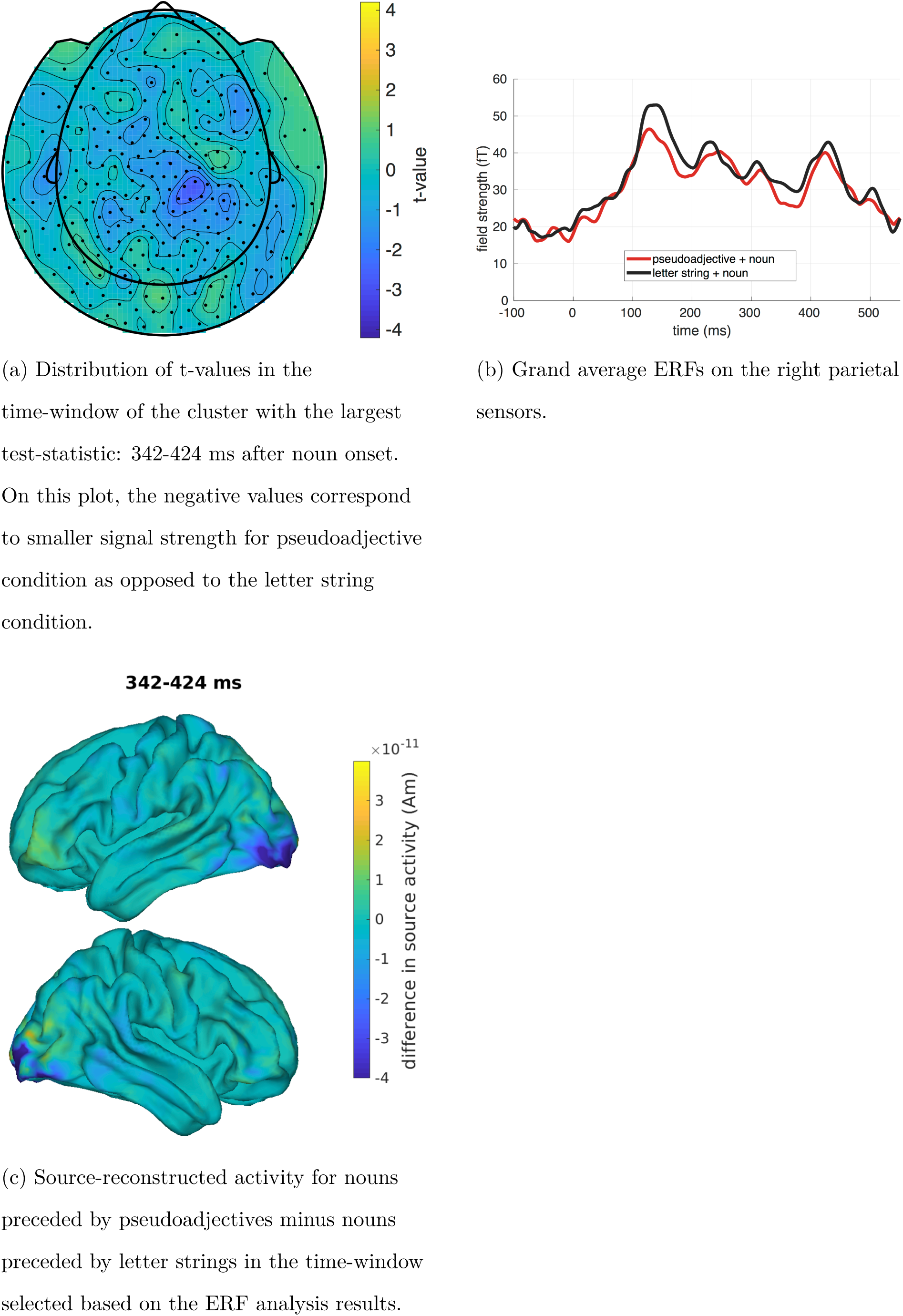
Plots with the results of the sensor-level analysis of syntactic composition: difference between nouns preceded by pseudoadjectives and by letter strings. Note that time-point 0 on these plots corresponds to the onset of the noun.

Overall, we conclude that there are no clearly interpretable results related to the composition effect we are interested in from these analyses of sensor-level data. This is not surprising given that previous MEG studies looking at the composition effect at the sensor-level also failed to show such effects.

## General discussion

We investigated how the brain carries out minimal adjective-noun phrase composition in Dutch, using magnetoencephalography. To investigate the semantic composition processes, we followed up on previous research, which contrasted processing of an adjective-noun phrase (using adjectives of different class) with processing of a noun preceded by a meaningless string of letters (Bemis & Pylkkänen, 2011; Ziegler & Pylkkänen, 2016). We adopted a methodology parallel to the previous studies as much as possible, but conducted our study in a different language and with additional norming criteria for the materials. For part of the research questions and the corresponding analyses, we had clearly specified hypotheses based on the results of previous studies. In addition, we also conducted exploratory analyses.

To investigate syntactic composition, we introduced a novel condition to the paradigm with pseudoadjectives instead of adjectives. In this condition, we expected syntactic composition to be carried out by the participants but not semantic composition. Since there was no previous MEG study that included such a condition, we relied on the results of fMRI studies to decide about the localization of the effects that we should observe and conducted exploratory analyses.

We failed to observe a composition effect, i.e., higher levels of activity when processing nouns preceded by a real adjectives as opposed to nouns preceded by letter strings. This was the case for a confirmatory analysis with a pre-defined time-window (i.e., 200-258 ms after noun onset) and region (i.e., BA21), and for the exploratory analyses (at any point between 100-500 ms after noun onset, in any of the four selected regions based on previous studies - BA21, BA38 [temporal pole], BA39 [angular gyrus], BA44+45 [left inferior frontal gyrus] - and for any type of adjective and noun). While absence of a difference in other regions is consistent with previous studies (e.g., Bemis & Pylkkänen, 2011; Blanco-Elorrieta et al., 2018; Blanco-Elorrieta & Pylkkänen, 2016 also do not find any semantic composition effect in the left BA44+BA45 or left BA39), absence of a difference in LATL (BA21 and BA38) is striking, given that multiple previous studies reported such a difference for a parallel contrast to the one we employed in our experiment (starting from Bemis & Pylkkänen, 2011, 2013a, 2013b; though see *Table 5* for a detailed overview of all findings and discussion below). Accordingly, we did not observe the previously reported modulation of activity in LATL by noun specificity and adjective class. These latter results are less surprising given that only two studies to date reported a modulation by noun specificity (Westerlund & Pylkkänen, 2014; Ziegler & Pylkkänen, 2016) and only one study reported a modulation by adjective class (Ziegler & Pylkkänen, 2016) in a similar set-up. Nonetheless, absence of the basic LATL composition effect precludes us from making conclusions about these *modulating* factors. We therefore do not discuss results regarding noun specificity and adjective class further. Finally, we also fail to observe a syntactic composition effect, i.e., higher levels of activity when processing nouns preceded by pseudoadjectives marked for grammatical gender as opposed to nouns preceded by letter strings.

**Table 5.**
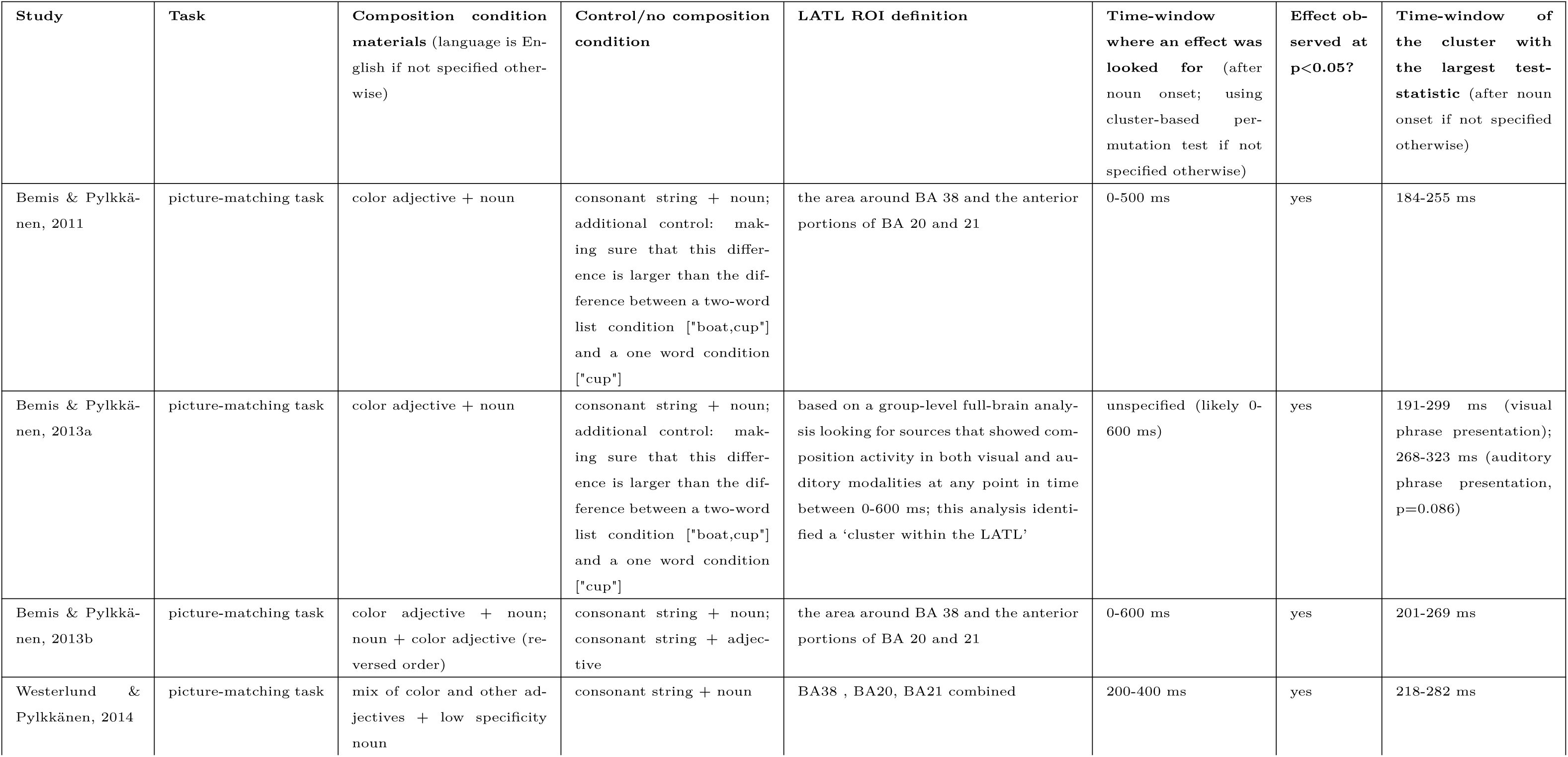
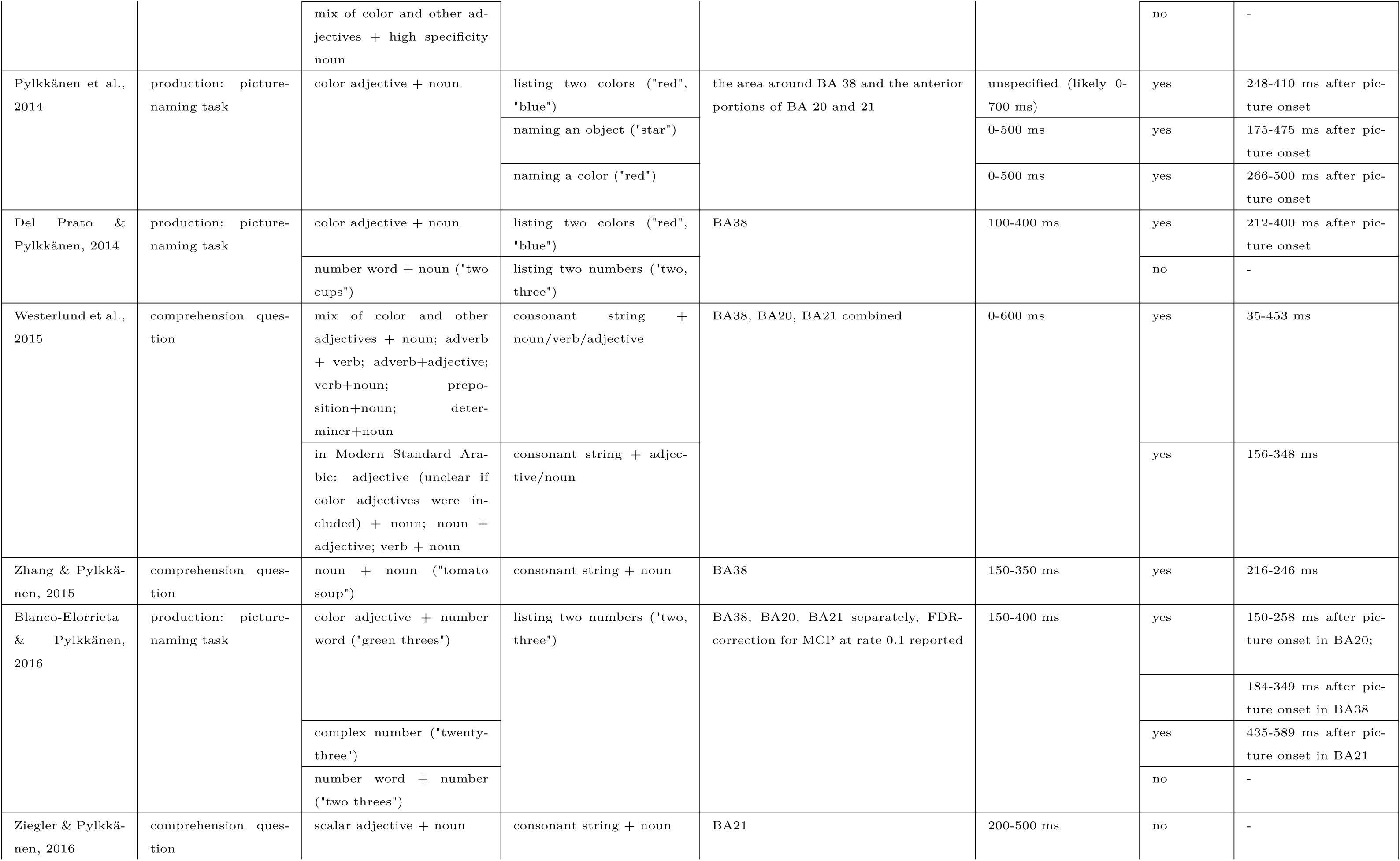
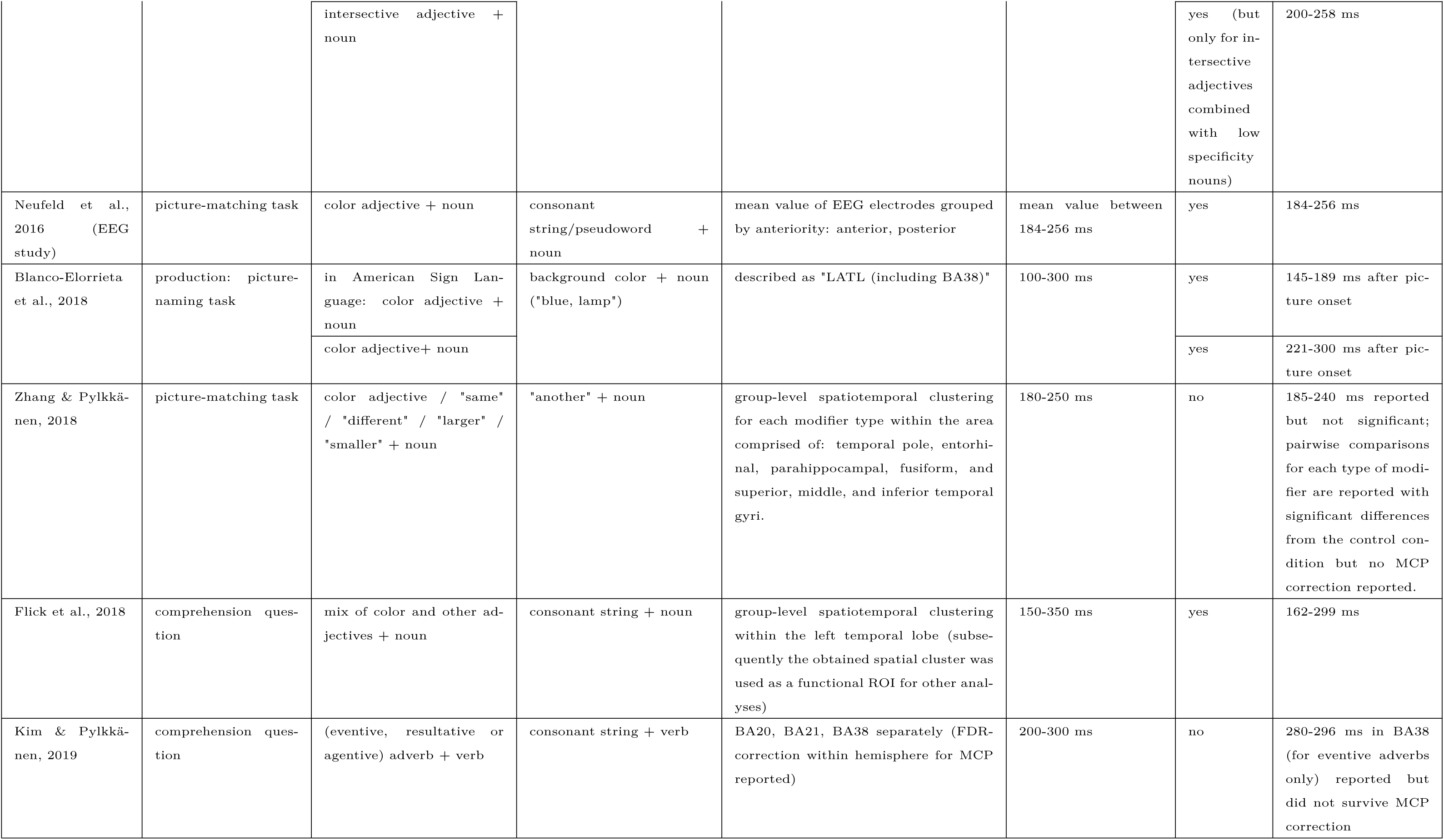
Summary of findings of studies that investigated composition of minimal two-word phrases using MEG. This table summarizes the results for LATL ROI analyses only. Only those comparisons are included in the table that are relevant to the presence/absence of the simple LATL composition effect (e.g., Flick et al., 2018 looked at compound nouns combined with adjectives which we omitted here).

Let us first discuss potential differences in data quality between ours and previous studies that could be responsible for failing to observe the previously reported LATL composition effect. Our study did not have a small number of participants or trials relative to the previous ones (our analyses included 33 participants, with 40 items per experimental design cell, whereas, for example, Z&P had 24 participants and 50 items per experimental design cell; other studies reported this effect with approximately 15-25 participants tested). Our MEG data artifact exclusion procedure is in parallel with or, perhaps, even improved in comparison to some of the previous studies (e.g., Bemis & Pylkkänen, 2011; Pylkkänen et al., 2014; Westerlund & Pylkkänen, 2014; Ziegler & Pylkkänen, 2016 etc. do not report removing heartbeat-related signal). In addition, whereas all previous studies to date (at least those we are aware of) demonstrating this effect used a scaled template brain for source activity reconstruction from MEG data, we used individual MRI scans of each participant which arguably resulted in more reliable localization of activity estimates. Finally and most importantly, in our sanity check analysis on the source reconstructed activity we replicated an independent previously reported effect; this means that the data of the present study was of adequate quality and that the pre-processing pipeline produced meaningful results. Given these considerations, we conclude that the signal-to-noise ratio in the present study was sufficient to detect the kinds of effects that have previously been shown. Therefore, it is unlikely to be the reason for the difference in the result from previous studies. Instead, we believe that content-related reasons (language, materials, task, analysis choices) should be considered.

### Potential reasons for the failure to observe the LATL composition effect

In this section, we discuss content-related differences between our study and the previous ones reporting the LATL composition effect that could have been the reason for our null effect. Such factors may warrant further investigation to probe the limits of and relevant preconditions for observing the LATL composition effect.

One potential reason why we did not observe an LATL composition effect could be that we conducted our study in Dutch, and that the properties of composition of minimal adjective-noun phrases in Dutch are such that an LATL composition effect might not show up in an MEG study. This is unlikely, however, given that Dutch is extremely similar to English, belonging to the same language family. The only notable difference between Dutch and English adjective-noun phrases is that in Dutch the adjectives are inflected for grammatical gender: an ending ‘e’ either is or is not added to the adjectives depending on whether they are combined with a noun belonging to one of two grammatical gender classes. This difference is unlikely to play a significant role since composition effects in left anterior temporal lobe in two-word phrases have also been reported for alternative phrase structures (specifically, for verb-noun, adverb-verb, adverb-adjective phrases in Westerlund et al., 2015, but see Kim & Pylkkänen, 2019 where only a weak effect was found and for just one class of adverbs in adverb-verb phrases) and in Arabic where adjectives also agree with nouns in terms of grammatical gender (Westerlund et al., 2015). Nonetheless, given that the evidence for the LATL composition effect is coming from only one previous study and with just one other language with grammatical gender agreement, there is a small likelihood that presence of grammatical gender agreement in Dutch could have resulted in the failure to observe the effect in the present study. If the absence of the effect in our study was indeed due to it being conducted in Dutch, it would mean that the presence of morphosyntactic agreement between an adjective and a noun somehow precludes involvement of LATL in composition. Alternatively, when morphosyntactic agreement is involved, the process is more heterogeneous than without such agreement, making it impossible to be detected. It is, for example, possible that the timing or the spatial extent of the composition process is less uniform across participants and/or nouns when morphosyntactic agreement processing is involved.

Another potential reason for the observed null effect might be a difference in stimulus materials and in the specific task we used when compared to previous studies. We reviewed all previous studies known to us that investigated the LATL composition effect, presented in *Table 5*. This table is not meant as an exhaustive review of all comparisons and contrasts reported in each study, but rather summarizes the results specifically for LATL ROIs with simple two-word phrases. Remember that the stimulus material of the present study consisted of scalar and intersective adjectives. When reviewing all previous studies, it is striking that, in fact, many of the studies reporting an effect used color adjectives only (7 out of 15) or color adjectives intermixed with other types of adjectives (another 2 out of 15). Only two studies reported the LATL composition effect without color adjectives in the materials (for information on the remaining four studies see footnote ^20^). One of these two studies was with noun-noun phrases rather than adjective-noun phrases (Zhang & Pylkkänen, 2015), with the other one being the Ziegler and Pylkkänen (2016) study that we followed in the present study. Since only two studies observed the LATL composition without color adjectives and one study (ours) does not find evidence for such effect, it is possible that the LATL composition effect is only robustly observed for adjective-noun phrases with color adjectives. If this is the case, it would mean that LATL is particularly sensitive to composition with color adjectives or that the composition process is more uniform across participants and/or nouns with color adjectives.

A similar line of reasoning is applicable to the specific task that participants had. In the present study, participants read a phrase and were asked to respond to a comprehension question at the end of each trial, with the question being about the combined adjective and noun meaning. As can also be seen in *Table 5*, the LATL composition effect has been reported predominantly in studies where participants matched a phrase to a picture or produced a phrase as a description of a picture. In contrast, only a small portion of the previous studies that observed the LATL composition effect reliably (4 out of 15) used a simple text-based comprehension task comparable to the task used in our study. Possibly, the presence of a picture in the task leads to participants being more likely to engage in imagery than a text-based comprehension task. Thus, we believe our null result then tentatively suggests that perhaps the LATL is more robustly involved in composition when participants have to engage in imagery of the objects^21^. The potential dependence of the LATL composition effect on the specific task used should therefore be investigated in future research. We stress that our proposal is deduced solely based on the pattern observed in the studies on minimal phrase composition, and in the future work it needs to be grounded in the wider literature on the functional role of LATL in language processing.

A striking aspect of previous studies concerns the variability in the regions of interest that were used in the analyses. As already noted in the *Introduction* section, while previous studies always referred to the region in which the composition effect was observed as the “left anterior temporal lobe”, its exact spatial extent did differ, sometimes dramatically, ranging from the anterior to the posterior portions of the left temporal lobe. Different studies defined the LATL as (all left) BA21, as BA38, as BA38+BA20+BA21 combined, as BA38+BA20+BA21 separately, as BA38 and the anterior portions of BA20 and BA21, by using spatial clustering in and around LATL or in the whole temporal lobe. The spatial extent of the observed effect is likely to differ somewhat in different studies given the limited spatial resolution provided by source reconstruction based on MEG data. However, given the breadth of definitions of the region of interest where an effect was reported, it is difficult to conclude with confidence that these are not, in fact, many different effects that were grouped under the same umbrella, or that at least some of these effects are not false positives^22^. To avoid these issues, in the present study we preferred to only conduct confirmatory analyses in exactly the same region of interest as the original study with the same design that reported the effect, with other regions being examined only in an exploratory way.

From a more general perspective, the present and previous studies assume that an identical cortical region (i.e., set of dipoles) across participants will perform an identical functional operation across participants – composition of an adjective and a noun. This assumption may not be warranted given that different (but probably neighbouring) cortical regions might be responsible for composition in different participants. This would in turn lower the strength of the signal in the averaged data (it should be noted that given the already low limited spatial resolution of MEG source reconstruction, small differences may not play a substantial role, but there would nonetheless be some reduction in the strength of the observed signal, depending on the extent of variability). An alternative and perhaps a better long-term solution for this line of research (solving the problem of researcher degrees of freedom with ROI definition described above and doing away with the assumption of identical cortical regions performing composition) would be to use a functional localizer for ROI definition. Such a functional localizer would use a pre-defined set of materials that would be identical across studies.^23^ and potentially localize the ROI on individual subject level (note that Flick et al., 2018 already used a functional localizer for ROI definition, but did so on a group level and with a novel set of materials; our suggestion to use an individual-level functional localizer is in fact rather ambitious given that many trials would need to be administered to each participant to reach a reasonable signal-to-noise ratio). Such a functional localization on an individual level before running group-level analyses has been previously proposed for fMRI studies (Fedorenko, Hsieh, Nieto-Castañón, Whitfield-Gabrieli, & Kanwisher, 2010; see also Matchin, Brodbeck, Hammerly, & Lau, 2019 which used fMRI data to define functionally relevant regions in each subject for the MEG analysis). In other words, the region of interest can be defined as the region showing increased activity for nouns preceded by real adjectives as compared to nouns preceded by letter strings for a particular participant given the same set of materials across studies. Whether different factors such as noun specificity and adjective class modulate the composition effect can then be investigated within this functionally defined ROI.

Overall, we conclude that our failure to observe the widely reported LATL composition effect is not as surprising after carefully considering the specifics of materials, tasks and regions of interest used in previous studies where such an effect *was* reported. Our review and discussion suggest that further investigation of limits and relevant preconditions for observing the LATL composition effect is needed.

### Syntactic composition effect

In order to investigate syntactic composition of adjective-noun phrases, we included a condition in which a pseudoadjective was combined with a noun. We expected that participants would carry out syntactic composition in these phrases without carrying out semantic composition since the pseudoadjective had an inflection that agreed with the grammatical gender of the noun. However, in principle our participants could also simply have ignored the pseudoadjectives and still respond correctly to the comprehension question (since it was only about the noun in this condition). As a way to check whether our participants did or did not pay attention to the pseudoadjectives, we added trials where the grammatical gender marking of the pseudoadjective and the grammatical gender of the noun did not match (i.e., syntactic violations). We expected to observe a grammatical violation signature in ERFs during processing of the noun in the form of an N400-like effect. Such an effect has been observed for grammatical gender agreement violations in adjective-noun phrases in other languages (Molinaro et al., 2011) and for grammatical gender agreement violations in determiner-noun phrases in Dutch (Hagoort, 2003). However, we did not observe such an effect, meaning that either participants did not perform syntactic composition in this condition or that we did not have enough power in this part of the study to observe a violation detection effect. The latter could indeed have been the case as we used only 20 trials with matching and 20 trials with mismatching pseudodjective inflection for this control analysis; we deliberately did so as we wanted to avoid fatiguing our participants with a longer experiment (note that this part of the study was administered at the very end of the experimental session). It is thus possible that the signal-to-noise ratio was not sufficiently high in this analysis for an effect to emerge. In a follow-up study, we would choose to administer more trials for this control analysis in order to be confident that we are not missing the violation detection signature. Another complication of our control trials concerns the fact that we could analyze data only up to 600 ms after noun onset (300 ms of the noun presentation followed by 300 ms of a blank screen after which a comprehension question was displayed). This trial structure was chosen in order to have it identical to the trial structure in the main part of the experiment on the LATL composition effect (and, therefore, to make sure that participants do not notice a difference and switch to a different mode of processing). But this implies that we cannot analyse the data of this control condition beyond 600 ms after noun onset and thus we cannot look at the time-window of a P600-like effect (approximately 500-800 ms after noun onset) which has sometimes been observed in addition or instead of an N400-like effect for similar grammatical violations (Hagoort & Brown, 1999; Molinaro et al., 2011). In a future study, it might be reasonable to have a longer blank screen after noun presentation in order to be able to look at the P600-like time-window as well.

Turning to the syntactic composition itself, we do not observe any syntactic composition effect, i.e., difference between processing nouns preceded by pseudoadjectives and nouns preceded by letter strings. This naturally leads one to the question of why we do not observe syntactic composition effects in our study, when such effects have been observed by others. It is possible that we do not observe such a difference because our participants simply did not perform syntactic composition (given the results with our control trials). Here, one potentially important difference between the present study and the fMRI study by Zaccarella and colleagues discussed above in the *Introduction* (Zaccarella & Friederici, 2015) is in the experimental manipulation: whereas in the materials of Zaccarella and colleagues real functional elements (determiners and prepositions) were combined with pseudowords as content words (e.g., ‘DIESE FLIRK’ [this flirk]), in our materials for testing syntactic composition there were no real functional elements. In fact, traditionally the fMRI research on syntactic structure building included real functional elements (see e.g., Matchin et al., 2017; Mazoyer et al., 1993; Pallier et al., 2011), and we do not know whether without these elements syntactic structure building proceeds in a comparable way. Therefore, it will be important for the future research to compare the more traditional set-up where minimal phrases include a real functional element and the present case where minimal phrases do not include a real functional element.

It is also possible that the participants *did* perform syntactic composition in our set-up, but we were not able to detect it with our method. As discussed in the *Introduction*, syntactic composition effects for minimal adjective-noun phrases have previously been reported in fMRI studies (Schell et al., 2017; Zaccarella & Friederici, 2015; Zaccarella et al., 2017), but there were no MEG studies specifically investigating it. It is possible that the time-course of syntactic composition is too variable between participants or phrases making it difficult to detect with averaged data with high temporal resolution as in the case of MEG, but possible with BOLD data which is capturing activity averaged over a longer time-span. Alternatively, spatial resolution of MEG source-reconstructed data is not good enough to capture the syntactic composition-related activity (for example, Zaccarella & Friederici, 2015 identify just a part of BA44 as most correlated with syntactic composition, a rather small patch of cortex). Note, however, that even with the spatial resolution of fMRI studies (and, related to the point raised in the previous paragraph, when real functional elements were included), syntactic composition effects have not always been observed for minimal phrases while they *were* observed for sentences (Matchin et al., 2017). In this context, some have suggested that syntactic composition of a simple minimal phrase like the one used here is highly automatic, hence not needing many resources to be processed and not detectable using our methods (suggested e.g., by Flick & Pylkkänen, 2018; Matchin et al., 2017; Pylkkänen, 2019; however, speaking against this possibility is the fact that some studies *do* observe syntactic composition effects in case of automatic syntactic processing, e.g., Hahne & Friederici, 1999; Hasting & Kotz, 2008; Pulvermüller & Assadollahi, 2007).

Another possible explanation for not observing syntactic composition effects in our study is that syntactic structure building may already have been initiated before the onset of the noun. Previous research showed that syntactic structure can be built in a top-down manner predictively - the language processing system may activate a certain syntactic structure before it has seen the syntactic class of the following items (e.g., Hale, 2001; Staub & Clifton Jr., 2006). In the minimal adjective-noun phrase paradigm that we used all items follow the same syntactic structure and, therefore, this structure is highly predictable. Hence, the syntactic structure could have been predicted already at the adjective position. A recent MEG study of phrases and sentences in blocked design supports a possibility of early syntactic structure predictions (Matchin et al., 2019). Future research will need to explore this possibility by manipulating the predictability of syntactic structure in the minimal phrase composition paradigm.

In summary, it remains unclear based on our findings whether ‘pure’ syntactic composition did not take place in our set-up or whether the methods we employed here are simply not appropriate for detecting such an effect. Moreover, given the absence of a basic composition effect there are no straightforward conclusions to be drawn about potential differences between composition involving semantics on the one hand, and ‘purely’ syntactic composition on the other.

## Conclusion

The present study investigated composition of minimal adjective-noun phrases in Dutch with two goals. The first goal was to look for a previously established composition effect in left anterior temporal lobe, as well as its modulation by noun specificity and adjective class; the latter modulation has been taken as evidence for the semantic nature of this composition effect. The second goal was to target specifically syntactic composition processes by including a novel condition where a pseudoadjective’s inflection matched the noun in terms of grammatical gender.

We failed to observe the previously reported LATL composition effect. Our review of previous studies that reported the LATL composition effect suggests that most likely this effect is only robustly observed when materials consist of color adjectives and/or include an imagery task. Thus, in our view, future research should focus on limits and relevant preconditions for observing the LATL composition effect. Additionally, our review reveals substantial inconsistencies in previous studies in terms of the brain regions and time-windows of interest that were used for analyses. We argue for more consistent definitions of regions and time-windows of interest in this line of research.

We did not observe a specifically syntactic composition effect either. However, because our control condition did not show with certainty that participants engaged in syntactic composition of pseudoadjective-noun phrases, we cannot be sure that the failure to observe this effect is not simply due to participants’ failure to engage in syntactic composition in this condition. We acknowledge that our control condition should have been statistically better powered, which we hope follow-up studies in a similar vein will ensure.

To conclude, the study of semantic (and syntactic) composition processes in minimal two-word phrases is surely a promising area of research as it allows one to study such composition processes in tightly controlled experimental settings. However, the present data also suggest that these tightly controlled settings should be used systematically to trace potential influences of the specific linguistic materials and tasks on composition effects before any general conclusions about semantic (and syntactic) composition can be drawn.

## Acknowledgments

We would like to thank the *Neurobiology of Language* group at MPI for Psycholinguistics/Donders Institute as well as the *Cognitive Semantics and Quantifiers* group at ILLC for fruitful discussions and their suggestions during early stages of the work on this project. We would also like to thank three anonymous reviewers for their thoughtful suggestions that helped us to improve the discussion of our results.

## Declarations of interest

None.

## Funding

This work was supported by the Netherlands Organization for Scientific Research (NWO) grant Language in Interaction [grant number 024.001.006].

## Footnotes

1 Here and later we mention the Brodmann Area to which the peak point of the observed difference in BOLD-signal belonged since in the present study analyses we define regions of interest (ROIs) in terms of Brodmann Areas.

2 Note that Schell and colleagues consider determiner-noun phrase composition ‘syntactically-driven’ whereas in the design of Matchin and colleagues such determiner-noun phrases are considered to have a semantic component (relative to the determiner-pseudoword condition). There is a large amount of theoretical and experimental work on whether determiners carry semantic information without a clear answer, so we do not take a stance or comment on this issue further, but simply note the discrepancy for clarity.

3 In fact, these adjectives are more commonly called ‘non-gradable’, with ‘intersective’ being only a subset of such adjectives (Kamp & Partee, 1995; Partee, 1995), but we will follow the terminology used in the Ziegler & Pylkkanen article for consistency. Note also that Ziegler & Pylkkanen (and accordingly the present study) do not take the strictly classical definition of ‘intersective’ adjectives. ‘Intersective’ is defined as not requiring a comparison class and operationalized using a ‘for a’ test; see below for details.

4 Note that to determine whether a participant should be included, we did not look at the number of remaining trials in the conditions in the additional data collection block of the experiment where pseudoadjectives with grammatical gender mismatch were presented (described below).

5 Z&P did not match nouns in their materials on frequency.

6 Note that we do not provide results of a test of significance of differences in terms of these prop-erties between conditions as is customary in language research since such a test has limited utility (see Sassenhagen & Alday, 2016).

7 Z&P also did not match adjectives in their materials on these features.

8 Z&P had 50 nouns in each noun specificity condition, and did not have a pseudoadjective condition, so their experiment had 300 trials in total.

9 Z&P do not report matching adjective-noun phrases in different conditions on plausibility. To assess whether a potential confound in plausibility of adjective-noun phrases used by Z&P could have been responsible for the observation of different activity levels for different adjectives classes, we also collected plausibility data for the set of materials used in the Z&P study. We collected data for this purpose in the same way as we did for our own materials, only translating the instructions from Dutch into English and recruiting native speakers of US English. We asked participants to give each phrase a score between 1 and 7 on how natural it sounds (more details, data collection and analysis scripts are available in the *Supplemental online materials*). Every adjective-noun combination received a score from 25 participants recruited in a web-based study. The intersective adjective-noun phrases in Z&P materials received a lower mean score - 5.67 (SD 1.03, range 3.08-6.96) - than the scalar adjective-noun phrases - 6.42 (SD 0.48, range 4.44-7; t(99)=-6.6, p<.001, d=-0.66). While adjective-noun phrases that we used also somewhat differ between conditions in terms of plausibility, the difference between them is substantially smaller (intersective adjective condition: mean 5.88 [SD 0.80, range 3.08-7]; scalar adjective condition: mean 6.08 [SD 0.68, range 3.48-6.69]; t[79]=-1.74, p=0.08, d=-0.19).

10 For comparison, Z&P report a mean score 2.54 for intersective adjectives and 5.44 for scalar adjectives.

11 We were restricted by the number of available participants in the pool of Dutch native speakers that we used for pre-tests.

12 Note that in Z&P the blocks were constructed manually such that no two adjectives or nouns were the same within each block, then the order of stimuli were randomized within blocks and the order of blocks was randomized; we preferred to instead fully randomize the order of stimuli

13 Z&P do not report dealing with heartbeat signal or muscle artifacts, but only report removing trials that were contaminated by eye movement artifacts.

14 Z&P used a template brain for source reconstruction.

15 Baseline correction is not mentioned in Z&P.

16 Note that Z&P used the Talairach atlas for separating activity into BA regions. Given that both atlases map Brodmann Areas, we do not expect that using a different atlas for parcellation will make a substantial difference in the observed results.

17 We constrained our time-windows to at least 100 ms after word onset since based on what we know about language processing, we do not expect to observe any effects at an earlier point; constraining the time-window this way allowed our analyses to be more focused and, therefore, more sensitive.

18 This is the criterion used in Z&P and other studies by Pylkkänen and colleagues. Note that we in addition ran analyses with a more common criterion p<0.05, and there were no substantial differences in the results.

19 For comparison, Z&P removed 24% of trials overall.

20 We are not certain that the materials in Standard Arabic used by Westerlund and Pylkkänen (2015) did not include color adjectives; the effect reported by Blanco-Elorrieta and Pylkkänen (2016) for complex numbers was much later than the expected LATL composition effect; finally, Zhang and Pylkkänen (2018) and (2019) report significant effects for other types of adjectives and for adverbs correspondingly, but in both cases only when analyzed separately and without multiple comparisons correction reported.

21 This, speculatively, is in line with the potential dependence on color adjectives being used since color adjectives are potentially more likely to involve imagery than other types of adjectives (compare: “white guitar” vs “large guitar”).

22 Note that this point of concern is also applicable to the differing time-windows that were analyzed for the presence of the effect across different studies.

23 One aspect to keep in mind with such functional localizers would be the potential differences in composition processes due to the morphosyntactic differences between languages (e.g., with and without morphosyntactic agreement between an adjective and a noun). This question would need to be investigated empirically.

## Notes

### Competing Interest Statement

The authors have declared no competing interest.

### Summary of Updates

Minor revision of the Discussion section; refining theoretical points made in the Introduction and Discussion sections.

